# Long term rescue of Alzheimer’s deficits *in vivo* by one-time gene-editing of *App* C-terminus

**DOI:** 10.1101/2024.06.08.598099

**Authors:** Brent D. Aulston, Kirstan Gimse, Hannah O. Bazick, Eniko A. Kramar, Donald P. Pizzo, Farbod Shavarebi, Trinity Rust, Sheila Garcia Rosa, Leonardo A. Parra-Rivas, Jichao Sun, Kristen Branes-Guerrero, Nidhi Checka, Neda Bagheri, Sara B. Rosenthal, Nihal Satyadev, Jared Carlson-Stevermer, Takashi Saito, Takaomi C. Saido, Anjon Audhya, Marcelo A. Wood, Mark J. Zylka, Krishanu Saha, Subhojit Roy

## Abstract

Gene-editing technologies promise to create a new class of therapeutics that can achieve permanent correction with a single intervention. Besides eliminating mutant alleles in familial disease, gene-editing can manipulate upstream pathophysiologic events and alter disease-course in wider patient populations. Here we use CRISPR-Cas9 to edit the last exon of amyloid precursor protein (*App*), relevant for Alzheimer’s disease (AD). Our strategy effectively eliminates an endocytic (YENPTY) motif at APP C-terminus in mouse and human neurons, while preserving N-terminus and compensatory APP-homologues. This manipulation favorably alters events along the amyloid-pathway – inhibiting toxic APP-β-cleavage fragments (including Aβ) and upregulating neuroprotective APP-α-cleavage. AAV-editing ameliorates neuropathologic, electrophysiologic, and behavioral deficits in an AD knockin mouse model. Effects persist for many months with no detectable abnormalities in germline-edited WT mice, and pathologic alterations in glial-transcriptomes are also normalized. Our strategy takes advantage of innate transcriptional rules that render terminal exons insensitive to nonsense-decay, and this upstream manipulation is expected to be effective for all forms of AD.

## INTRODUCTION

The ability to treat diseases at a genetic level with a single intervention and bring about a permanent cure is poised to transform the practice of medicine ^1^. Recent trials in hematologic and systemic disorders have reported unprecedented clinical outcomes ^2, 3^, leading to the first approvals of CRISPR therapies ^4^; raising hopes that similar strategies can be employed for neurodegenerative diseases like AD where traditional therapeutics have been largely disappointing ^5^. Although recent trials using monoclonal antibodies to Aβ assemblies show efficacy ^6^, clinical benefits are modest and the drugs need to be repeatedly administered throughout life, increasing risks of intracranial hemorrhage ^6, 7^. Moreover, many studies have shown that a number of APP-β-cleavage products – other than Aβ – can also cause pathologic deficits such as endo-lysosomal dysfunction ^8–10^, and removing extracellular Aβ would not be effective against these toxic intermediates. With the promise of targeting etiology and achieving “permanent correction” ^11^, gene-based therapies offer an alternative.

Recent studies have begun to explore gene-editing in AD. CRISPR-based strategies have been used to selectively inactivate disease-associated mutant alleles in AD, while keeping WT alleles intact ^12–14^. Such allele-specific inactivation reverses biochemical abnormalities in cells ^13^, and one study has shown that AAV-mediated delivery of CRISPR-components by local injections can ameliorate some deficits in a transgenic (“5x familial AD”) mouse model ^14^. Besides caveats of CRISPR-editing in over-expression-based models where unknown transgene-copies are expressed, at best, these strategies would only be effective in a small number of familial AD patients with known gene mutations [< 1% ^15^]. In fact, few mutation-independent CRISPR-based therapeutic approaches have ever been reported ^16^, and such interventions are expected to be beneficial for large patient populations. Local brain injections of nanoparticles carrying CRISPR-Cas9 have also been used to inactivate β-site APP-cleaving enzyme 1 (BACE1) in mice ^17^, but there is broad consensus that BACE1 is not a good clinical target ^18^. Moreover, local brain injection is unlikely to be effective in AD, given the widespread pathology. An exploration of CRISPR-based germline *App*-edits in mice revealed a 3’UTR-deletion that decreased *App* expression ^19^, but safety and therapeutic implications have not been evaluated.

Our translational efforts have centered around APP, that has an established role in AD pathogenesis ^20, 21^. Remarkably, a single amino-acid *APP* variant (*A673T* ‘Icelandic mutation’) is protective for sporadic AD ^22^, underlining the importance of this gene in both familial and sporadic cases. One therapeutic approach is to attenuate the entire *APP* gene, but *APP* (or *APP*-like) genes are highly conserved across species, implying an important physiologic role ^23^. Moreover, protective APP-cleavage products – that are physiologically expressed – would also be eliminated in this scenario. While deposition of APP-β/γ-cleavage products is a hallmark of AD, under normal conditions, APP is largely cleaved by an alternative α-cleavage pathway, generating distinct APP-fragments that are neuroprotective and neuroregenerative ^24^. APP cleavage by α-secretases preclude β-cleavage ^20, 24^, and in principle, this shift in the balance of cleavage (β/g ◊ α) can be therapeutically manipulated for both sporadic and familial AD. Interestingly, the protective Icelandic *APP* mutation is also thought to make APP a less favorable substrate for β-cleavage and Aβ production ^22, 25^. Our current studies follow up on previous work from us and others, showing that the YENPTY-motif at the APP C-terminus is critical in mediating the trafficking of APP into endosomes enriched in BACE1 and triggering APP/BACE1 interaction, which is the rate-limiting step initiating the β/γ-cleavage pathway ^26–32^. More recently, we showed that elimination of the APP C-terminus containing the YENPTY motif attenuated β-cleavage and augmented α-cleavage ^29^. Though our previous proof-of-principle experiments showed that this approach can work in cells, effects on AD pathology and behavior are unknown, and safety of this approach has not been examined *in vivo*. Importantly, translational goals – such as a therapeutically viable strategy for widespread brain delivery or therapeutically relevant means of editing human *APP* – remain unrealized.

## RESULTS

Our approach is based on CRISPR-Cas9 editing by non-homologous end joining (NHEJ), which is the most studied of all editing approaches, and also the basis of almost all clinically-relevant applications to date ^2, 3, 33^. A conceptual schematic demonstrating altered trafficking of APP after C-terminus deletion is shown in **Figure 1A**. Note that loss of the YENPTY-motif (post-editing) blocks internalization of surface APP into endosomes containing BACE1, attenuating APP-β-cleavage. Consequently, APP-α-cleavage is augmented – presumably due to increased dwelling of edited APP on membranes where α-cleavage is predominant ^34^ – leading to a shift of APP cleavage pattern upon gene-editing. In conventional applications, NHEJ leads to premature stop-codons and nonsense decay of the entire transcript, resulting in a loss of the protein ^35^. However, our strategy takes advantage of innate transcriptional rules that prevent nonsense decay when premature termination-codons are installed in the last exon and ∼ 50-55 nucleotides upstream of the last exon-exon junction – reviewed in ^36^ (also see discussion). Note that the YENPTY motif and flanking regions within the APP C-terminus are encoded by the last exon (exon 18) of *APP* (**Fig. 1B**).

**Figure 1:**
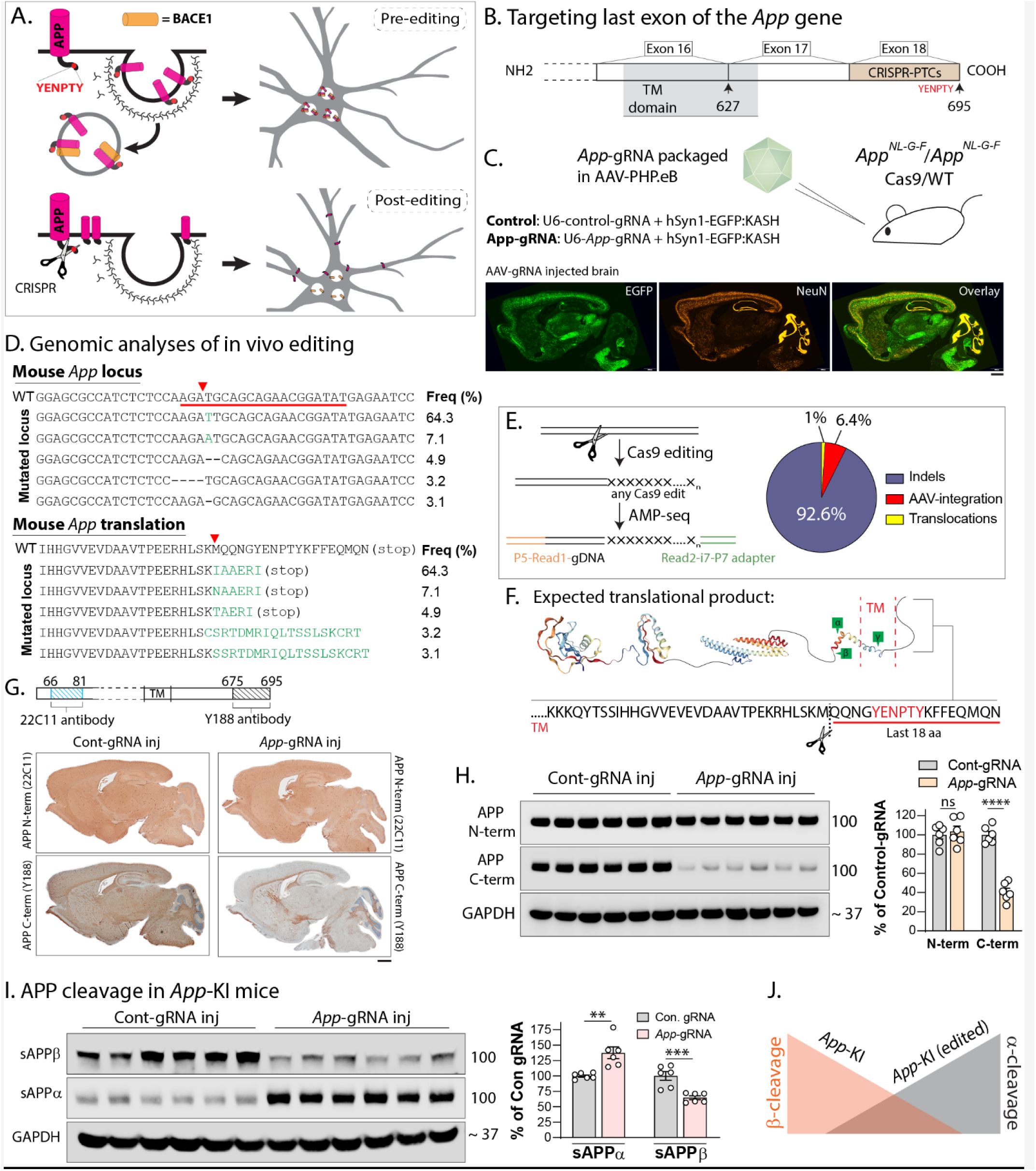
AAV-CRISPR-editing of *App* C-terminus favorably alters APP β/α cleavage pattern *in vivo*. **A)** Schematic showing internalization of APP into endosomes containing BACE1 (top). CRISPR-mediated effective deletion of the APP-YENPTY motif (scissors, below – see results for details) prevents the endocytosis of APP and its interaction with BACE1, which is the rate-limiting step triggering APP-β-cleavage pathway (schematic attribution: Alec Nabb). **B)** Note that the YENPTY motif lies within the region encoded by the last exon (exon 18) of APP, and CRISPR-induced premature termination codons (PTCs) installed within the last exon are expected to generate a truncated protein (see results and discussion for details, TM = transmembrane domain). **C)** A construct including a gRNA targeting the last *App* exon – along with an EGFP marker – was packaged into the AAV-PHP.eB capsid and intravenously injected into *App^NL-G-F^*/Cas9-KI mouse brains. Representative images of AAV-injected mouse brains (below) showing broad transduction (also see **Supp.** Fig. 1A). All data here were obtained from mice injected at 1.5 months and evaluated at 12 months. All western blots were from cortex and hippocampus. Scale bar = 1 mm. **D)** Major mutated *App* loci resulting from above *in vivo* CRISPR-editing, and their relative frequencies (top), with expected translational products (bottom). Note that most of the editing outcomes predict short indels. **E)** Analysis of editing outcomes using quantitative AMP-seq (left) to identify insertion/deletion (indel), translocation, or AAV-integration at Cas9-cleavage sites. Quantification of editing outcomes – expressed as percentages of total edits – in AAV/*App*-gRNA injected mice. Note that indels are the most common outcome of editing, with minimal off-target effects (also see **Extended data Fig. 1B-F**). **F)** Schematic showing segment of APP (last 18 amino-acids, containing the YENPTY motif) that is expected to be eliminated after CRISPR-editing. **G)** Schematic on top shows recognition sites for APP N-and C-terminus antibodies (22C11 and Y188 antibodies respectively). Note that upon editing, The Y188 antibody is not able to recognize the translational product, which is used as a surrogate marker for editing in our experiments. Immunostains of mouse brains (bottom panels) show selective attenuation of Y188-signal in *App*-gRNA injected mice (also see **Supp. Figs. 2A-B**). Scale bar = 1 mm. **H)** Western blots from mouse brains (cortical/hippocampal lysates, JH49 antibody)) also show selective attenuation of the Y188-antibody signal, quantified on right (N=6 mice per condition – see full blots in **Supp.** Fig. 2C). **I)** Western blots from soluble *App^NL-G-F^*/Cas9-KI mouse brain fractions (cortical/hippocampal lysates) to evaluate β/α cleavage products (see full blots in **Supp.** Fig. 2D). Note reversal of the β/α cleavage pattern in *App*-gRNA injected mice; quantified on right (N=6 mice per condition). **J)** Conceptual schematic showing changes in β/α cleavage after *App*-editing. All quantitative data presented as mean +/- SEM; ns=non-significant, ** p<0.01, ***p<0.001, ****p<0.0001 (unpaired t-test).

### AAV-driven *App* C-terminus editing favorably alters the balance of APP cleavage

To edit *App in vivo*, we packaged a *App*-gRNA (“676-gRNA”, numbering based on predicted amino-acid cut-site and brain isoform APP-695) – or control-gRNA not targeting to any known sequence (**Extended data Table 1**) – and a fluorescent marker into AAV-PHP.eB, which is an engineered capsid that allows widespread transduction into mouse brains after intravenous injections ^37^. This AAV-cargo was delivered into homozygous *App*-KI mice crossed with Cas9-KI mice, leading to broad transduction of mouse brains (**Fig. 1C**; also see **Extended data Fig. 1A** and **Extended data Table 1**). The *App*-KI mice in our experiments have a humanized Aβ domain, carry three familial AD mutations (Swedish, Arctic, and Iberian – *App^NL-G-F^*), and accrue Aβ plaques and other deficits over time ^38^. Unlike most other AD mouse models, the *App^NL-G-F^* mice do not overexpress APP, and the gene is driven and regulated by native transcriptional and translational elements. Notably, APP overexpression is known to induce pathologic changes in various model-systems ^39, 40^, and generates abnormal APP fragments that can confound interpretation ^41^ – concerns that are not applicable in the *App*-KI mice.

First we asked if our AAV-CRISPR injections edited the mouse *App* gene as expected. Genomic analysis of the AAV injected *App*-KI mice showed that edits occurred in the expected *App* loci and led to premature stop codons within the last exon, with relatively short insertions and deletions (indels, **Fig. 1D**). Quantification of editing outcomes in these samples using amplicon-sequencing [AMP-seq ^42^] showed that indels were the most common outcome at the cut-site, followed by AAV-integration and translocations, without detectable off-target effects (**Fig. 1E**, also see **Extended data Fig. 1B-F**). Note that the last 18 amino-acids – including the pentapeptide YENPTY motif – are expected to be deleted by our editing strategy (**Fig. 1F**). To verify that our *App* editing generated a translational product that was truncated at the C-terminus, we used an antibody (clone Y188) that only recognizes the last 20 amino-acids of APP, combining this with antibodies against APP N-terminus (**Fig. 1G**, top). While full-length APP is recognized by the Y188 antibody, if the APP C-terminus is missing, this antibody will not recognize the truncated protein [see further characterization of these antibodies in our previous study^29^]. As shown in the mouse brain section in **Figure 1G** – top panels, the N-terminus APP antibody showed widespread staining of APP in both control- and *App*-676-gRNA injected brains. However, Y188 staining was selectively attenuated in *App*-676-gRNA injected mice, compared to mice injected with the control-gRNA (**Fig. 1G** – bottom panels). Two-color immunofluorescence showed that most of the AAV-transduced neurons (∼ 90%) also had attenuated Y188-antibody staining, suggesting efficient editing (**Extended data Fig. 2A**). Western blots using N- and C-terminus antibodies also confirmed the selective C-terminal truncation of APP. While signals from the Y188 antibody were significantly attenuated in *App* edited samples, band-intensities in western blots using an APP N-terminus antibody were unchanged (**Fig. 1H**, see full blots in **Extended data Fig. 2B**).

Next, we asked if editing the *App* C-terminus led to a shift in the balance of APP β/α cleavage *in vivo*. Cleavage of APP by β/α secretases lead to secreted APP-cleavage products (sAPPβ/α) that can be biochemically detected in soluble brain fractions. As expected, brains from *App*-KI mice with control-gRNA injections had a substantial increase in sAPPβ, compared to sAPPα (**Fig. 1I** – left lanes). However, brains of *App*-KI mice injected with the AAV-CRISPR payload had a marked reduction in β-cleavage products, with an increase in α-cleavage fragments (**Fig. 1I** – right lanes, quantified in **Fig. 1J**; see full blots in **Extended data Fig. 2C**). Overall, the data indicate that somatic editing of the *App* C-terminus using AAV vectors led to a shift in the APP-β/α cleavage pattern (**Fig. 1K** – also see our previous study ^29^ for further characterization of β/α cleavage post-editing in mouse and human cells). Note that these changes in β/α cleavage are unlikely to be due to alterations in levels of the secretases ADAM10 (A Disintegrin And Metalloprotease domain-containing protein 10) and BACE1, which are not significantly affected by our editing strategy (**Extended data Fig. 2D**).

### AAV-driven *App* C-terminus editing rescues multiple deficits in *App*-KI mice

Next we asked if altering the balance of β/α APP-cleavage using our approach also attenuated pathologic phenotypes in the *App*-KI mice. Schematic in **Figure 2A** shows the experimental plan of AAV injections and evaluation timepoints. First, we injected mice just before the emergence of Aβ pathology in this model [∼ 2 months ^38^]. As shown in exemplary images (**Fig. 2B**), Aβ plaques were attenuated after *App*-676-gRNA injection. Evaluation of Aβ pathology over the course of our experiments showed that while plaque deposition continued in animals injected with the control-gRNA, the Aβ-attenuating effect of the *App*-676-gRNA was additive over time, likely due to ongoing dampening of APP β-cleavage for several months after a single injection (**Fig. 2C**, also see **Extended data Fig. 2E**). Substantial reductions in insoluble Aβ were also seen in the brains of one-year old animals (**Fig. 2D**). The reduction in Aβ pathology, and increase in neuro-protective sAPP α-cleavage fragments was seen as early as 2.5 months after injection (**Extended data Fig. 2F-G**). Endo-lysosomal pathology is a prominent feature of AD ^10^, and such abnormalities in the *App*-KI mice were also diminished by App-676-gRNA injections (**Fig. 2E** – quantified on right).

**Figure 2:**
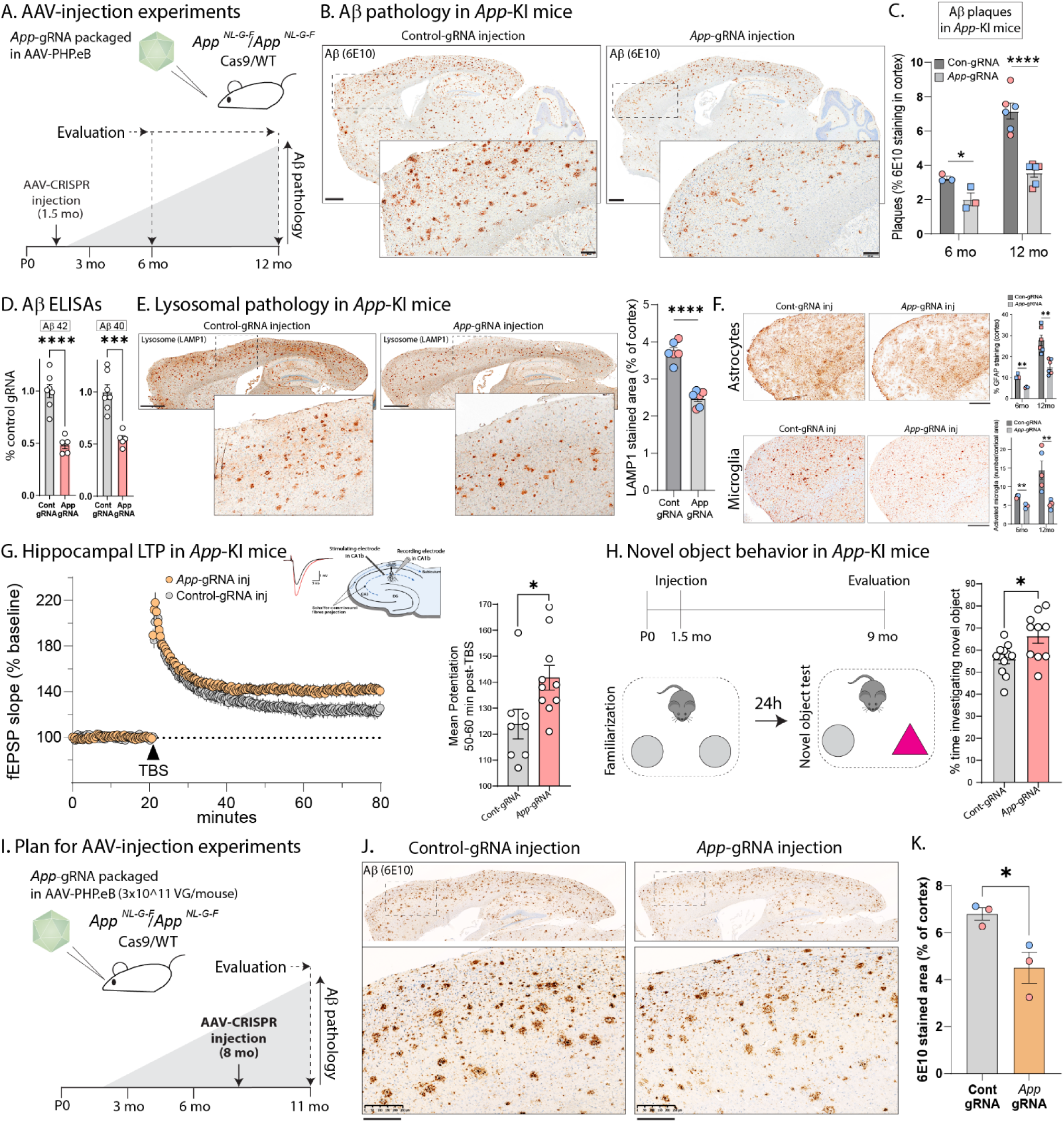
AAV-driven CRISPR editing of the *App* C-terminus ameliorates multiple deficits in *App*-KI mice. **A)** Schematic showing timepoints of AAV-*App*-gRNA injections and evaluation of neuropathology in *App*-KI/Cas9-KI mice. **B)** Representative sections showing Aβ pathology in one-year old mice, following protocol in **A)**. Note decreased Aβ plaques in *App*-gRNA injected animals (zoomed frontal cortex). Scale bars: main = 1 mm, zoomed inset = 200 μm. **C)** Quantification of Aβ progression. While Aβ deposits increase over time in control-gRNA injected brains as expected, there is a marked attenuation in *App*-gRNA injected animals (N=3-6 mice per condition;– datapoints: blue = male, red = female). **D)** Aβ ELISAs from GuHCl soluble brain fractions (cortex and hippocampus) in one-year old mice. Note marked Aβ attenuation in *App*-gRNA injected mice (N=5-7 animals for each condition). **E)** Representative sections showing lysosomal pathology in one-year old *App*-KI mice. Note attenuated lysosomal staining in *App*-gRNA injected animals, quantified on right (N=6 animals for each condition). Scale bars = 1 mm. **F)** Representative frontal cortical sections showing staining of astrocytes (top) and microglia (bottom) in *App*-KI mice injected with control-gRNA or *App*-gRNA, with quantification on right. Note sustained attenuation of glial pathology after a single *App*-gRNA injection (N=3-6 mice per condition; also see **Extended data Fig. 2K-M** – datapoints: blue = male, red = female ). Scale bars = 500 μm. **G)** LTP recordings from acute hippocampal slices of *App*-KI mice (10-month-old) injected with control or *App*-gRNA at 1.5 months; data quantified on right. Note augmentation in LTP after *App*-gRNA injections. (N=8-10 slices per condition– also see **Extended data Fig. 3A**). **H)** Novel object test behavior in 9-month-old old mice (schematic on top). Note that the *App*-gRNA injected mice spend more time exploring the novel object, compared to controls. (N=10-11 animals per condition). All quantitative data presented as mean +/- SEM. **I)** AAV-*App*-gRNA injections after onset of pathology in 8 month-old *App*-KI mice. Note decreases in Aβ (**J-K**), as well as microglial pathology (**Extended data Fig. 3B-D**; N=3 animals for each condition). All quantitative data presented as mean +/- SEM; *p<0.05, ** p<0.01, ****p<0.0001 (unpaired t-test).

Recent studies have revealed an important role of neuroinflammation in the progression of AD pathology. Neuroinflammatory markers are increased in AD, and human genetics have uncovered several AD-risk genes that have functions in innate immunity ^43^. Innate immune responses in the brain are primarily mediated by glia, and an increase in neuroinflammation has also been documented in *App^NL-G-F^* mice ^41^. Astrocytosis and microglial-activation is also attenuated in the *App*-KI mice after a single *App*-676-gRNA injection, and these attenuating effects were also additive over time (**Figs. 2F** and **Extended data Fig. 2H)**. Specific populations of disease-associated microglia (DAM) are thought to play a role in AD ^44^. Indeed, immunohistochemical staining for TREM2 – a key marker of DAM – was attenuated after *App* editing, though there was no significant change in the homeostatic microglial marker P2RY12 (**Extended data Fig. 2I**; see **Extended data Fig. 2J** for further analyses of glial-plaque interactions). Previous studies have shown that long-term potentiation (LTP) is impaired in acute hippocampal slices from older *App^NL-G-F^* mice ^45^. LTP reflects the persistent strengthening of synapses in response to recent activity and is thought to be the cellular mechanism underlying learning and memory. Thus, LTP impairments in the *App^NL-G-F^* mice are especially relevant in the context of a memory disorder such as AD. We compared LTP in acute hippocampal slices from 10-month-old *App^NL-G-F^*/Cas9 mice that were injected with AAV-*App*-676-gRNA (or control-gRNA) at 1.5 months. As shown in **Figure 2G**, there was an augmentation of LTP in the *App*-676-gRNA injected group compared to the control-gRNA injected group (editing in slices was verified post-hoc, see **Extended data Fig. 3A**). Impairments in the novel-object recognition test have also been reported in older *App^NL-G-F^*mice ^46^. Novel-object recognition is a measure of innate exploratory behavior in the absence of external cues, and based on the logic that cognitively intact mice show a natural preference for new objects in their environment. Previous studies have shown that older *App^NL-G-F^* mice have deficits in this test, showing a decreased preference for novel objects ^46^. Accordingly, we performed this test in 9-month-old *App^NL^ ^G-F^*/Cas9 mice that were injected with AAVs carrying *App*-676-gRNA (or control-gRNA) at 1.5 months. As shown in **Figure 2H**, the *App*-676-gRNA injected mice spent significantly more time exploring the novel object compared to controls, suggesting rescue of cognitive function upon *App* editing. Rescue of Aβ and microglial pathology was also seen when these AAVs were injected in 8 month-old *App^NL-G-F^*/Cas9 mice – well after the onset of pathology – though astrocytosis persisted (**Fig. 2I-K** and **Extended data Fig. 3B-D**). Comparable results were obtained with another gRNA that edited *App* slightly upstream of the 676-position, but still within the last exon (*App*-659-gRNA, see **Extended data Fig. 3E-L**.

### Germline editing of the *App* last-exon in *App*-KI and WT mice

To evaluate the safety of our gene-editing approach, we generated mice where the last-exon of *App* was edited in the germline of WT mice. We used the *App*-659-gRNA in these experiments, reasoning that since these germline-edited mice lacked the C-terminus of APP from birth – with the 659-gRNA editing a larger segment of the APP C-terminus – examining older mice from this group would be a stringent test of safety. Accordingly, we injected *App-*659-gRNA/Cas9 ribonucleoprotein complexes into zygotes from WT mice (and also *App*-KI mice, see next), and selected two germline-edited strains with indels within the last exon, expected to effectively delete ∼ 35 amino acids from the APP C-terminus throughout the brain (**Extended data Fig. 4A-B** - named “Δ5” and “insert-T” based on generated indels. Western blots of brain homogenates were consistent with a selective C-terminal truncation of APP in genomically edited mice (**Extended data Fig. 4C**). Importantly, our *App*-editing strategy had no effect on the *App* homologues APLP1/2 due gRNA target-sequence mismatch (**Extended data Fig. 4D**). Note that APLP1/2 are known to compensate for loss of *App* function in mice, and also contain YENPTY domains [reviewed in ^47^]. Interestingly, WT *App* editing also led to a decrease in endogenous APP β-cleavage and an increase in α-cleavage (**Extended data Fig. 4E**), suggesting that our CRISPR-driven manipulation of β/α cleavage can favorably alter cleavage pattern of WT APP *in vivo* – relevant in the context of sporadic AD without *APP* mutations. For further details on genotyping strategies, see **Extended data Fig. 4F-H** and Methods. Examination of brains from WT germline-edited mice did not show any gross deficits in neurons, synapses, astrocytes and microglia, when compared to their WT counterparts (**Extended data Fig. 5A-C**), and both groups performed similarly in memory tests (**Extended data Fig. 5D**).

Next we tested our germline-editing approach in *App*-KI mice, generating *App*-KI-Δ5/InsT mice that lacked the APP C-terminus from birth (**Fig. 3A**). As expected, there was a marked suppression of APP-β-cleavage and augmentation of APP-α-cleavage in germline-edited *App*-KI mice (**Fig. 3B**). In all experiments so far, we used homozygous (*App^NL-G-F^*/*App^NL-G-F^*) mice, mainly because the relatively faster development of pathology in these animals is more amenable to experimentation. To test our editing strategy in a more plausible clinical scenario, we performed germline *App* editing in heterozygous (*App^NL-G-F^*/WT) mice and evaluated Aβ pathology in older (10.5-month) animals. Strikingly, Aβ plaques were almost absent in the genomically deleted *App^NL-G-F^*/WT mouse strains (**Fig. 3C** and **Extended data Fig. 5E**). Since almost all familial AD patients are heterozygous, and the magnitude of neuropathology is generally comparable in familial and sporadic AD brains, these data suggest that *APP* C-terminus editing at the earliest possible stages in AD may potentially have profound therapeutic effects.

**Figure 3:**
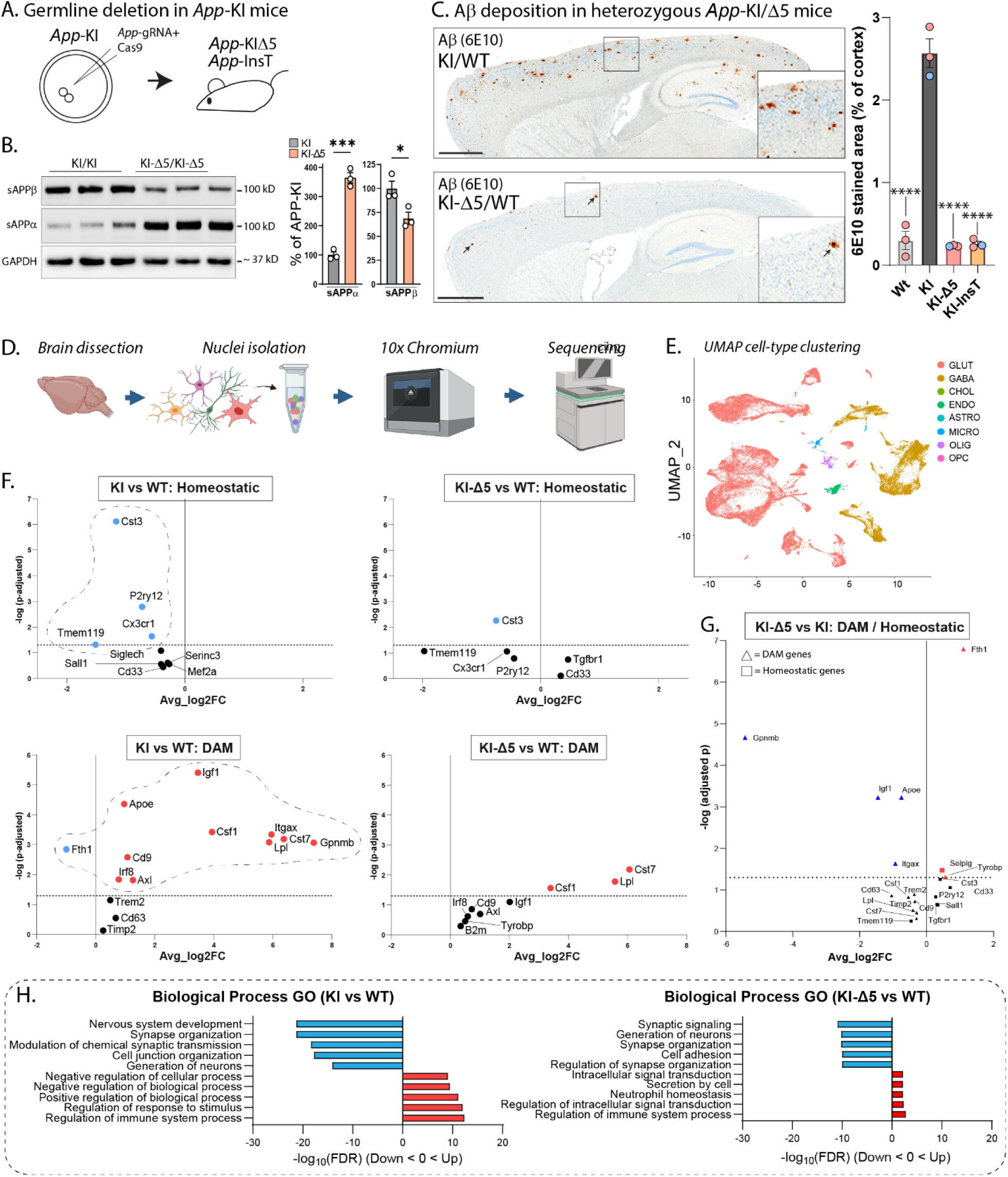
Germline *App* C-terminus editing in *App*-KI and WT mice. **A)** Strategy for generating *App*-KIΔ5/InsT mice by germline CRISPR-deletion. Zygotes from *App*-KI mice were injected with *App*-gRNA/Cas9 ribonucleoprotein complexes targeting the *App* C-terminus, and resulting founders were screened and outbred to produce stable heterozygous and homozygous lines (also see **Extended data Fig. 4**). **B)** Reversal of APP β/α cleavage pattern in germline-edited *App*-KI mice, quantified on right (N=3 animals/condition, cortical/hippocampal lysates). **C)** Representative sections of Aβ staining in germline-edited heterozygous *App*-KI mice. Note that Aβ plaques are almost absent in the edited mice (bottom panel, small arrowheads point to rare plaques, also see **Extended data Fig. 5E**), quantified on right. Scale bar = 1mm, data shown as mean +/- SEM. N = 3/condition. * p < 0.05, ***p < 0.001, **** p < 0.0001 and ns = non-significant. **D)** Workflow for sNuc-Seq from *App*-KI and *App*-KIΔ5 mouse brains (N = 2 brains per genotype). Transcriptomic profiles of isolated nuclei were generated with 10x chromium sNucSeq, followed by deep sequencing of cDNA libraries. **E)** UMAP (Uniform Manifold Approximation and Projection) visualization (right) shows efficient sorting into different cell-types (also see **Extended data Fig. 6C**). **F)** Volcano plots of homeostatic microglia (top panels) and disease associated microglia (DAM, bottom panels) in *App*-KI or *App*-KI-Δ5 mice, compared to WT mice (differentially expressed genes marked by dashed ovals). Note that most of the dysregulated genes in the *App*-KI mice are normalized in the *App*-KI-Δ5 mice. **G)** Ratio of DAM/homeostatic genes in *App*-KI or *App*-KI-Δ5 mice show that core markers of the DAM signature (*Apoe*, *Gpnmb*, *Igf1*, *Itgax*) are downregulated after gene-editing. Significance was defined as an adjusted p-value of <0 .05 in all Snuc-seq experiments. **H)** Biological Process gene-ontology (GO) analysis of all differentially expressed genes (DEGs) in microglia from *App*-KI or *App*-KI-Δ5 mice, compared to WT mice. Bar graphs indicate top five significantly down-regulated (blue) or up-regulated (red) biological pathways. All quantitative data presented as mean +/- SEM; *p<0.05, ***p<0.001, ****p<0.0001 (unpaired t-test for Fig. 3B, one-way Anova for Fig. 3C).

### *App* C-terminus editing rescues microglial transcriptomic changes in *App*-KI mice

Neuroinflammation and microglial activation have emerged as important players in the pathologic progression of AD ^43^. Besides human genetics, the relevance of microglia in AD pathology has been further bolstered by studies using single-cell/nuclei sequencing that can overcome limitations posed by cell-type heterogeneity and diversity of disease phenotypes. Transcriptomic studies of microglia in AD mouse models show gradual transitions from a homeostatic to a disease-linked state, revealing a set of disease associated microglial (DAM) genes that are altered with disease progression ^44^, which are also seen in the *App^NL-G-F^* knockin model ^48^. Accordingly, we asked if our *App* editing strategy also ameliorated some of these disease-linked alterations in microglia. To ensure a homogenous background of edited cells, we used the homozygous *App*-KI germline-deletion (*App*-KI-Δ5) mice for these experiments. As expected, glial activation was significantly attenuated in 10-month-old *App*-KI mice with germline-deletion of the C-terminus (**Extended data Fig. 6A-B**), along with Aβ plaques and associated pathologies. The experimental flow of the sNuc-Seq experiments is shown in **Figure 3D**. Briefly, ∼ 19,000-23,000 high-quality nuclei from 10-month-old mouse brains were analyzed to compare cellular-molecular maps between *App-*KI and *App*-KI-Δ5 animals. The nuclei were partitioned into multiple cell-types following standard protocols ^49^ (**Fig. 3E**, also see **Extended data Fig. 6C**), and sNuc-Seq data from *App*-KI-Δ5/*App*-KI mouse brains were compared to brains from WT animals.

For data analysis, our two-step goal was to first identify glial genes that were differentially altered in the *App*-KI brains compared to WT brains, and then ask if these alterations were abrogated in the *App*-KI-Δ5 animals. Additionally, we also compared our sNuc-Seq data to published RNA-seq databases from AD mouse models^48^. Expectedly, homeostatic genes were downregulated, while DAM genes were upregulated in *App*-KI mice (**Fig. 3F** – left panels, dysregulated genes highlighted by dashed ovals). Strikingly, most of these dysregulated microglial genes are normalized in the *App*-KI-Δ5 animals (**Fig. 3F** – right panels, **Fig. 3G**, and **Extended data Fig. 7A-B**). **Figure 3H** shows the top ten dysregulated pathways in KI v/s WT and KI v/s KI-Δ5, and additional transcriptomic data from these animals is shown in **Extended data Figures 7C-G**. Overall, the data show that many transcriptomic changes in *App*-KI mice are ameliorated by C-terminus editing.

### Amelioration Alzheimer’s pathology by a single AAV-vector carrying all CRISPR-components

Towards a therapeutically viable option for delivering our gene-editing payload, we developed a single AAV-vector carrying all CRISPR-components necessary for editing the C-terminus of *App*. The overall design of the vector and the targeted genomic sequence is shown in **Figure 4A**. Considering the ∼4.7 kb limit of AAV packaging, we expressed SaCas9 – a compact ^33^ and clinically-relevant ^33, 50^ genome editor – via a recently described neuron-specific promoter *pCALM1*, which is based on the human *CALM1* gene ^51^. At 120 bp, *pCALM1* is only about 1/5 the size of the human synapsin-1 promoter (∼470-500 bp) that is typically considered for CNS gene therapies ^52, 53^, and this strategy allowed us to insert the full WPRE element in the expression construct, while staying within the 4.7 kb packaging limit. The WPRE element has an established role in boosting AAV gene expression ^54^, and this design is expected to lower the therapeutic dose, potentially augmenting the safety of future AAV therapies (also see **Extended data Fig. 8B**). A 3xHA-tag was also added to identify transduced neurons, and we chose a PAM-site/genomic-target that was close to the *App*-676-gRNA used in our previous AAV-injection experiments. The single-vector AAV-CRISPR payload was intravenously injected into homozygous *App*-KI mice at ∼1.5 months, and brains were evaluated at 4 months for transduction efficiency and *App* C-terminus editing (using Y188 immunoreactivity as a surrogate marker of editing as described previously). Un-injected and control-gRNA injected mice were used as controls. Note widespread transduction of neurons in the AAV-injected brains (HA staining) including hippocampal CA1/2 regions, along with loss of Y188 signal and relative preservation of the APP N-terminus (**Fig. 4B-C**). As expected, *App* editing led to favorable alteration of APP-β/α cleavage (**Extended data Fig. 8C**), along with reduced Aβ plaque pathology and diminished insoluble Aβ as evaluated by ELISAs (**Fig. 4D-E**). Lysosomal and microglial pathology was also attenuated in the *App*-KI mice after AAV-gRNA/SaCas9 injections (**Fig. 4F-G**, also see **Extended data Fig. 8D-F**), although there was an increase in astrocytosis (**Fig. 4G**), perhaps due to AAV-driven inflammation.

**Figure 4:**
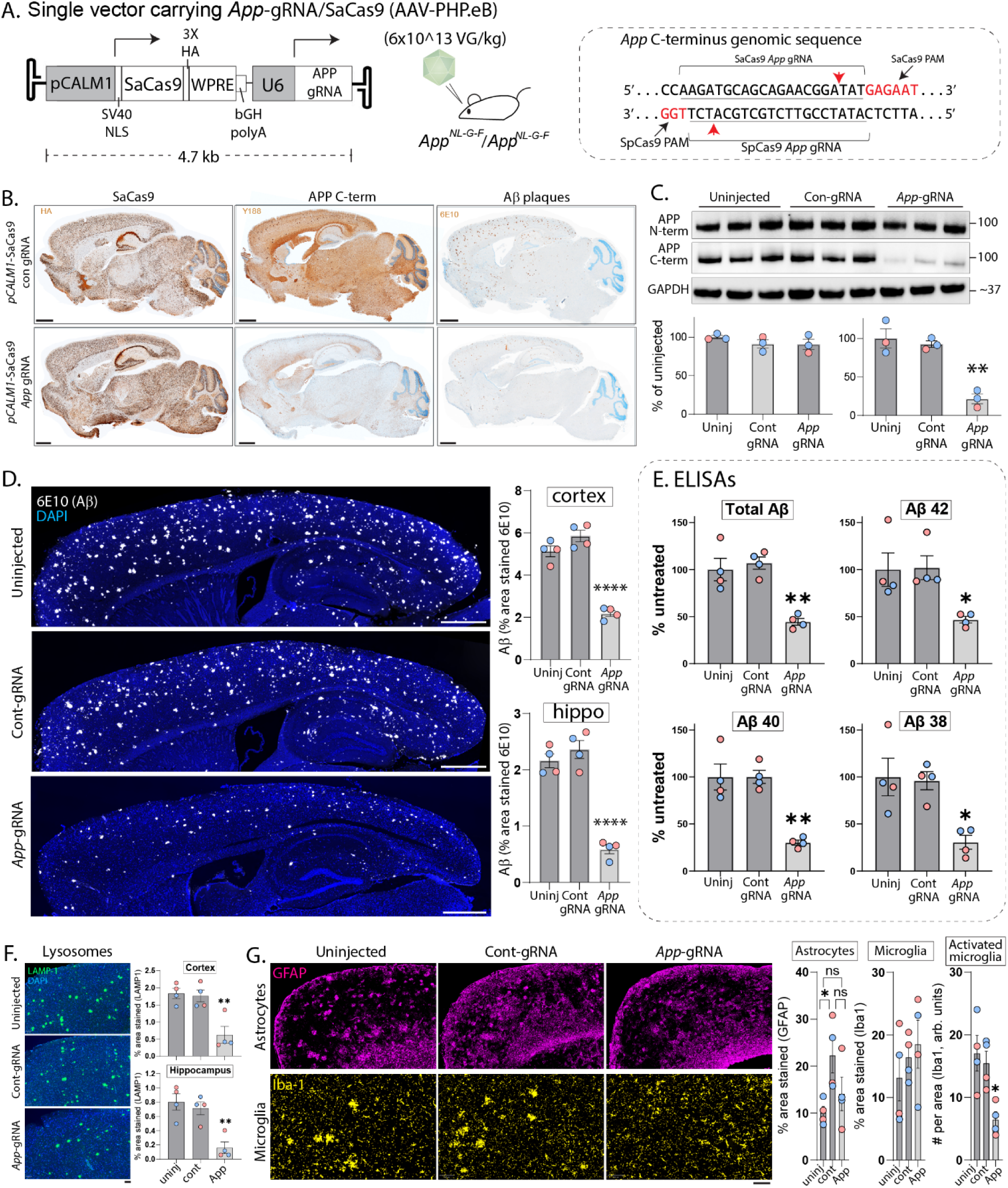
A single AAV carrying all CRISPR components attenuates neuropathology in *App*-KI mice. **A)** Schematic showing vector design accommodating the ultra-compact *pCALM1* promoter, *App*-gRNA driven by the U6 promoter, and a HA-tag (to identify transduced neurons) between the AAV-ITRs. *App*-KI mice (homozygous) were intravenously injected with the single AAV carrying all CRISPR-components at 1.5 months, and brains were evaluated at 4 months (uninjected and control-gRNA injected mice were used as controls). Note that the position of the SaCas9 PAM/cut-site (red arrowheads) in the *App* C-terminus are close to the SpCas9 cut-site (right). **B)** Adjacent brain sections stained with HA (to identify transduced neurons), Y188 (proxy marker of edited neurons), and 6E10 (Aβ) antibodies. Note broad transduction in all cases with attenuation of Y188 and 6E10 staining in *App*-gRNA injected mice (scale bars = 1 mm). **C)** Western blots from cortical/hippocampal lysates show attenuation of signal with the Y188 (APP C-term) antibody, with relative preservation of the N-term in *App*-gRNA injected mice (also see **Extended data Fig. 8C**; N=3 mice/condition – datapoints: blue = male, red = female). **D)** Representative Aβ stained sections of control and *App*-gRNA/SaCas9 injected mouse brains. Note attenuated Aβ staining in the *App*-gRNA-injected brain (scale bars: main = 800 μm; N=4 mice/condition – datapoints: blue = male, red = female). **E)** Aβ ELISAs from brains (GuHCl soluble fractions) show marked Aβ reduction in *App*-gRNA injected brains, compared to controls (N=4 mice/condition – datapoints: blue = male, red = female). **F-G)** Representative frontal cortical sections showing attenuation of lysosomal and microglial pathology, but with some astrocytosis in *App*-gRNA-injected brains (also see “results”; N=4 mice/condition – datapoints: blue = male, red = female). Scale bars = 500 μm. All data presented as mean +/- SEM; *p<0.05, **p<0.01, ****p<0.0001 (one-way ANOVA).

Finally, we looked for a human-relevant gRNA/Cas combination that would 1) effectively edit *APP* and truncate the YENPTY domain in human APP, and 2) fit into the 4.7 kb packaging limit of AAVs. Towards this we identified 7 gRNAs targeting a suitable window within the last exon of *APP* that were also compatible with the compact and AAV-adaptable SaCas9 (or a variant called KKH SaCas9 ^55^); see **Figure 5A**. These gRNA/Cas combinations were tested in human iPSC-derived neurons using a lentiviruses/antibiotic-selection based strategy that generates a near-homogenous edited population and avoids confusion due to mixture of edited and un-edited neurons (see schematic in **Fig. 5B** and Methods). AMP-seq data from edited neurons showed that while all 7 gRNas effectively edited *APP*, YENPTY-disruption was highest with 5 of these gRNAs (also see **Extended data Fig. 8G-H**). ELISAs confirmed Aβ reductions with these 5 gRNAs (**Fig. 5E**), suggesting that the single-vector AAV strategy using a compact Cas and human-relevant gRNA targeting the last exon of *APP* could be potentially effective in humans.

**Figure 5:**
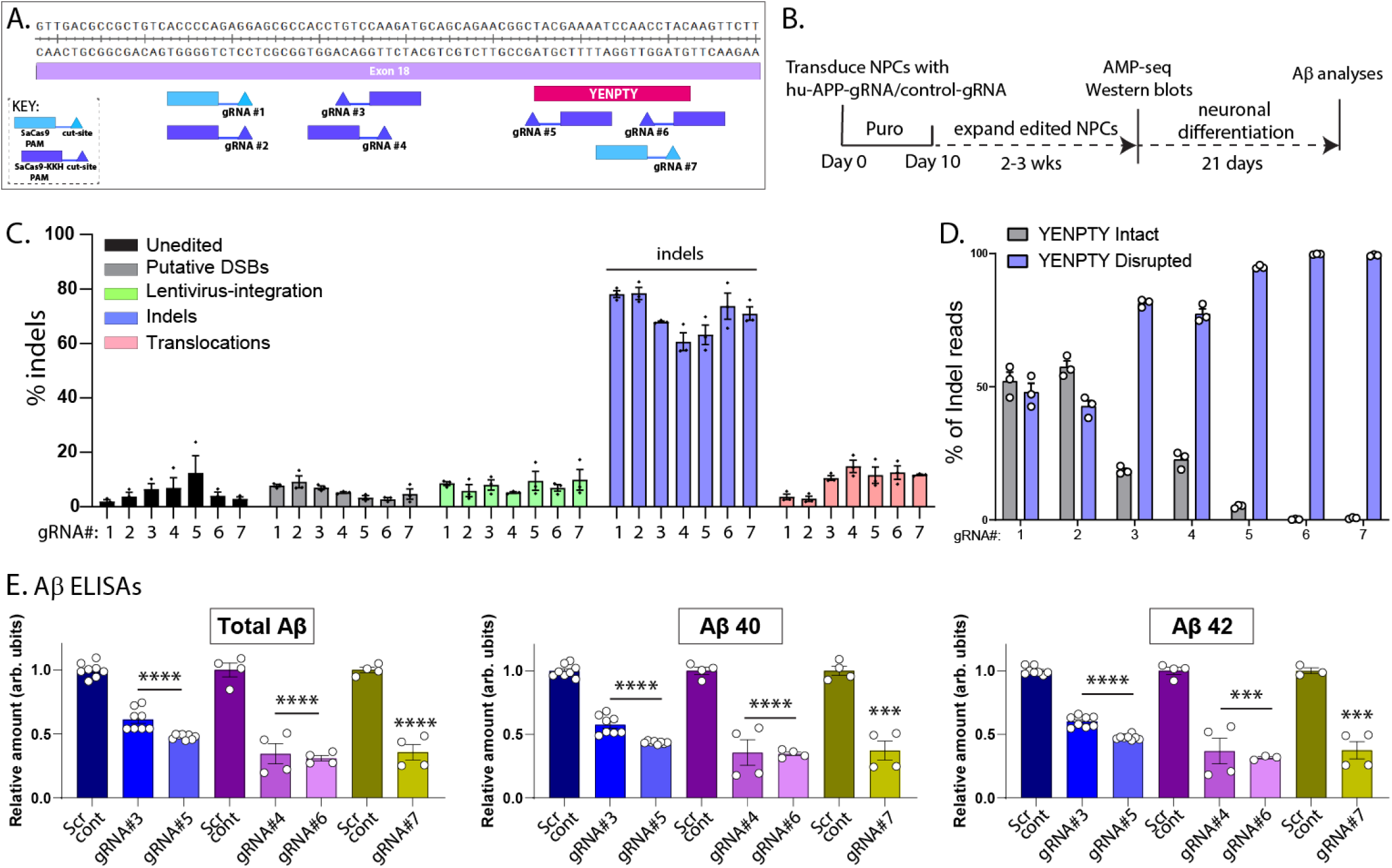
Human *APP* C-terminus editing with compact and AAV-adaptable Cas enzymes. **A)** Schematic showing PAM/cut-sites of 7 gRNAs targeting the last exon (exon 18) of human *APP*, YENPTY region is marked with a red box. Note that only PAM sites compatible with compact AAV-adaptable Cas enzymes were considered - SaCas9 (5’-NNGRRT-3’ PAM) and SaCas9-KKH (5’-NNNRRT-3’ PAM). Also note that gRNAs #1 and #2 have the same PAM/gRNA sequence that was paired with either SaCas9 or SaCas9-KKH. **B)** Experimental design for generating an edited population of human neurons. Human neural progenitor cells (NPCs) were transduced with gRNA/Cas9 using lentiviruses, cells carrying the CRISPR payload were selected using puromycin, and differentiated into neurons. AMP-seq and western blots were done in edited NPCs and media from differentiated neurons were used for Aβ ELISAs. **C, D)** Summary of editing outcomes from AMP-seq (N = 3 independent transductions for each Cas/gRNA pair). Note that all gRNA/Cas combinations led to indels **(C)**, with gRNA#3-7 showing the greatest disruption of YENPTY **(D)**. Black bars in **(C)** represent % of AMP-seq reads that deviated from WT *APP* sequence. Grey bars represent the % of reads with YENPTY disruption, calculated by subtracting the number of unedited/unproductive edits (reads with edits that did not disrupt the YENPTY motif, see **Extended data Fig. 8H**) from the total number of reads. NPC lines with maximal YENPTY disruption (gRNA#3-7) were selected for neuronal differentiation and Aβ analyses. **E)** Aβ analyses from conditioned media of edited neurons shows that gRNAs 3-7 also attenuate Aβ. Data represents mean amounts (relative to scrambled controls) +/- SD. N = 3-8 wells (2 independent transductions) for all Cas9-gRNA pairs. ****p <0.0001, ***p < 0.001 (one-way ANOVA).

## DISCUSSION

Here we demonstrate a CRISPR-Cas9 based therapeutic strategy to effectively delete a pentapeptide endocytic motif at the C-terminus of APP and favorably alter the balance of APP cleavage – attenuating toxic APP-β-cleavage, while upregulating neuroprotective APP-α-cleavage. This manipulation rescues neuropathologic, electrophysiologic, behavioral, and transcriptomic changes in an AD knockin mouse model, and a single injection of AAVs carrying CRISPR-components rescued deficits for many months. Early germline editing of the *App* C-terminus in a more AD-relevant heterozygous *App*-KI background essentially eliminated Aβ pathology, suggesting that when combined with early intervention, this approach may have substantial therapeutic effects. Similar germline editing in WT mice had no detectable anatomic or behavioral consequences *in vivo*, attesting to the safety of this approach. Finally, our experiments in human neurons show that AAV-compatible Cas/gRNA combinations can edit the C-terminus of human *APP*, effectively truncate the YENPTY domain, and lead to decreased Aβ secretion. Collectively, our studies offer a path for developing a one-time disease-modifying therapeutic for AD.

### Incidental features in the *APP* gene that favor our editing strategy

Indels generated by CRISPR-Cas mediated NHEJ usually lead to premature stop codons in the DNA ^35^. Typically, eukaryotic cells see these synthetic termination signals – proximal to the native stop codon – as abnormal, and consequently, innate repair mechanisms are triggered to eliminate the mRNA via nonsense-mediated decay, leading to a loss of the protein. However, recognition of a stop codon as ‘abnormal’ depends on the position of the termination codon relative to the last exon-exon junction, and extensive previous work has shown that premature stop codons within the last exon do not lead to decay of the transcript [reviewed in ^36^]. Briefly, transcriptional rules dictate that if a premature termination signal lies within the last exon – or within ∼50 nucleotides upstream of the last exon-exon junction – innate mechanisms do not recognize this transcript as abnormal, and instead, translation is terminated at the premature stop signal, which is expected to generate a truncated protein. Fortunately, about half of the APP C-terminus, including the YENPTY-domain and flanking regions, is encoded by the last exon (exon 18), allowing us to manipulate APP in a way that preserves the transmembrane-domain and N-terminus, permitting physiologic α-cleavage and generating soluble APPα fragments. The precision of CRISPR-editing also allows normal production of APP-like proteins [APLP1 and APLP2, see **Extended data Fig. 4D** and ^29^]. APLP1/2 are encoded by separate genes, and also have YENPTY-domains ^47^, and it is expected that protein-protein interactions mediated by this domain will continue to occur post-editing. Extensive in-vivo studies in mice have also established that *Aplp1*/*2* can compensate for *App* function ^47^, and the lack of any detectable abnormality in WT *App* C-terminus edited mice (**Extended data Figs. 4-5**) may also be attributable to this fortuitous scenario.

### A potential one-time gene editing therapeutic for all forms of Alzheimer’s disease

In principle, our upstream manipulation of APP-cleavage is expected to be therapeutic for sporadic AD, as well as familial AD with *APP* and presenilin (*PSEN*) mutations. The etiology of sporadic AD is unclear, and likely complex. While studies suggest that there is deficient Aβ clearance in AD patients ^56^, human genetics also implicate upstream genes in the amyloid pathway like *APP, PSEN1/2* and *ADAM10* ^15, 57^, and BACE1 activity is also reportedly higher in sporadic AD brains ^58^ – incriminating Aβ overproduction. Regardless, our upstream manipulation to tilt APP cleavage towards a more physiologic state is expected to be effective for sporadic AD, and we have previously shown that our strategy can favorably alter APP α/β-cleavage and attenuate Aβ in WT human neurons *(29)*. However, in the absence of a reliable model for sporadic AD, efficacy cannot be directly tested without clinical trials. Notably, a shift in APP processing towards the non-amyloidogenic pathway is also thought to be the basis for neuroprotection in the *APP A673T* Icelandic mutation, seen in sporadic AD population ^22, 25^, further suggesting that a similar approach could work in sporadic AD. Studies have also shown that secreted α-cleavage products sAPPα – but not sAPPβ – can rescue hippocampal LTP deficits in APP knockout mice ^59^, implying that a sustained increase in sAPPα may be effective in AD. Regarding timing of treatment, while earliest possible intervention is obviously the best (and a universal medical principle for any therapeutic), it is encouraging that our approach ameliorates pathology in *App*-KI mice, even when administered months after the onset of symptoms (**Fig. 2I-K**).

Our previous studies demonstrated the efficacy of our approach in isogenic human iPSC-lines carrying the *APP* London mutation ^29^. Taken together with the data shown here in the *App*-KI mice (efficacy in a mouse model with humanized Aβ-domain and three familial AD *APP*-mutations), the collective evidence strongly suggests that our approach would be effective for familial AD with *APP* mutations. Regarding familial AD with *PSEN* mutations, it is thought that the increased Aβ aggregation is due to the relative abundance of longer Aβ peptides that are more aggregation-prone ^60^. More recent studies suggest that *PSEN* mutations may lead to a loss of function in processing APP-β-CTF (C-terminal fragments) ^61^. Regardless, since our approach attenuates APP-β-CTF fragments that are required for subsequent cleavage by the presenilin/γ-secretase complex, it is expected that our strategy would be therapeutic in this scenario. However, there are ∼ 300 presenilin mutations in familial AD, and further focused work is needed to test therapeutic efficacy and safety of *APP* C-terminus editing in this setting. Widespread delivery into larger primate brains remains a challenge for all gene-therapy approaches targeting the brain. Nevertheless, several ongoing clinical trials are relying on intracisternal injection of AAVs into the cerebrospinal fluid for broad transduction (NCT04747431, NCT04408625, NCT03634007, NCT04127578), and emerging technologies are expected to offer better solutions in the near future ^62, 63^ .

### Limitations of the Study

As mentioned above, definitive proof of efficacy and safety in sporadic AD can only come from human trials, and broad transduction of a large brain is also challenging, which are general caveats relevant for many applications of gene-based therapies for neurodegenerative diseases. Also, focused efforts will be necessary to validate our gene-editing approach in the setting of various *PSEN* mutations found in familial AD patients, as noted above. High AAV doses can be toxic ^64^, and we saw astrocytosis in the doses used for single-vector AAV experiments, though we did not see any gross anomalies. Though our doses are comparable to the human doses used in AAV9-based Zolgensma gene therapy for spinal muscular atrophy ^65^, therapeutic approaches that require widespread delivery would greatly benefit from further technical improvements in genome editing and vector development.

In summary, our studies demonstrate the potential of a one-time CRISPR-Cas9 based gene-editing treatment for AD. Though further work is needed for human translation, our vision is to develop a universal ‘one-and-done’ gene therapy for AD. Potentially, our gene-editing therapeutic could also be used as a combination-therapy in elderly patients; delivered after initial doses of amyloid-attenuating agents (such as lecanemab) and abrogating the need for frequent intravenous injections throughout life that substantially increases risks of intracerebral hemorrhage.

## Methods

### List of antibodies used in this study

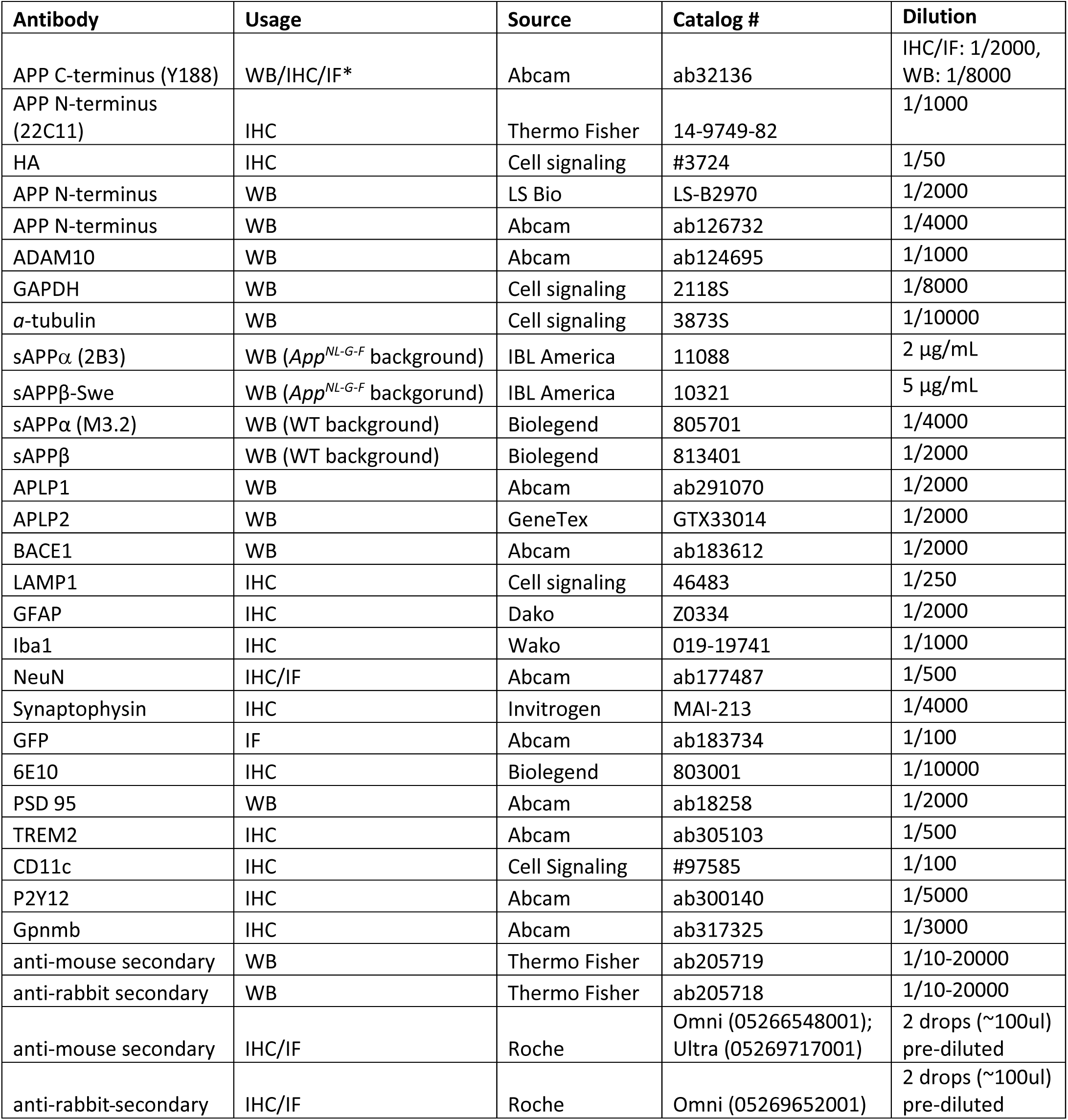

### Mouse breeding and CRISPR deletion of *App* C-terminus in vivo

#### General procedures

All animal procedures were done according to guidelines set by the University of California, San Diego. Mice were euthanized by CO_2_ asphyxiation and the brains were rapidly dissected out. Tissue was snap-frozen on dry ice (for ELISAs/biochemistry) or drop-fixed in 4% paraformaldehyde (for histology).

#### Generation of App^NL-G-F^/Cas9 knockin mice

*App^NL-G-F^*(*App*-KI) mice harboring a humanized Aβ domain and three familial AD mutations – Swedish, Arctic and Iberian^38^ – were obtained from RIKEN Brain Science Institute (Japan). KI mice were then crossed with CAG-driven SpCas9 knockin mice from Jackson labs (stock # 028239^66^) to produce *App*-KI/SpCas9 KI mice. Mice used for AAV-driven SpCas9/*App*-gRNA injection experiments were homozygous for the *App* knock-in and hemizygous for SpCas9 knockin. Mice used for SaCas9/*App*-gRNA (single-vector) experiments were only homozygous for the *App* knock-in allele. For AAV-injection experiments, littermates (both males and females) were randomly selected for transduction with AAVs carrying scrambled control-gRNA (not targeting to any known region in the mouse genome) or *App*-gRNA. For genotyping of the *App*-KI/SpCas9 KI mice, genomic DNA was isolated from ear clips using QuickExtract buffer (Lucigen) following manufacturer’s instructions. Knock-in alleles were then PCR amplified from genomic DNA templates using RedExtract PCR mix (Sigma), and amplicons were visualized on agarose gels with SybrSafe (Thermo Fisher).

#### Generation of App genomic-deletion mice

These mice were generated at the University of Wisconsin-Madison Genome Editing and Animal Models Core. One-cell C57BL/6J or *App*-KI fertilized embryos were microinjected with 50 ng/μl *App* C-terminus targeting guide-RNA and 40 ng/μl Cas9 protein (PNA Bio, Newbury Park, California). Injected embryos were transplanted into pseudo-pregnant B6D2 F1 females. Pups were sequenced at weaning to screen for gene-edits. Two APP C-terminus indels, a T base pair insertion (insT) and a 5bp deletion (Δ5), were selected during screening of founders and selected for colony propagation. Founders were crossed with C57BL/6J mates and F1 mice were sequenced. F1 WT-Δ5/InsT or *App*-KI-Δ5/InsT mice were then crossed with C57BL/6J mice for an additional 2 generations. Finally, F3 generation mice were crossed to produce homozygous WT-Δ5/InsT or *App*-KI-Δ5/InsT mice. For genotyping of Δ5 mice, genomic DNA was isolated from ear clips using QuickExtract buffer (Lucigen) following manufacturer’s instructions. Genomic region containing the CRISPR-edit was PCR-amplified using OneTaq polymerases in Standard Buffer (New England Biolabs). After PCR, 1μl of AciI restriction enzyme was added to 50µl of PCR reaction sample and samples were incubated at 37°C for 1 hr. DNA bands were then visualized agarose gels with SybrSafe (Thermo Fisher).

### AAV vectors – cloning, production, and intravenous injections

#### Cloning, production and purification of AAV vectors

For experiments in *App*-KI/Cas9 mice, scrambled control or *App*-targeting gRNAs were cloned into pAAV9-U6sgRNA(SapI)_hSyn-GFP-KASH-bGH vectors at Sap1 site as previously described^29^. SaCas9 AAV vector and wild-type SaCas9-3XHA coding sequences were derived from pX601, Addgene plasmid #61591 (a gift from Feng Zhang). *pCALM1* promoter sequence was derived from Addgene plasmid #213973 (a gift from Zhonghua Lu, Shenzhen Institutes of Advanced Technology). Lentiviral vector and WPRE sequences were derived from either lentiCRISPR v2, Addgene plasmid #52961 (a gift from Feng Zhang, MIT), or pCK002, Addgene plasmid #85452 (a gift from Aviv Regev, Genentech). To generate SaCas9 KKH coding sequence (following ref # ^55^), mutagenic primers were used with the Q5^®^ Site-Directed Mutagenesis Kit (New England Biolabs, E0554) to generate amino acid missense mutations E782K, N968K, and R1015H in wild-type SaCas9-3XHA from pX601. To generate single vector pCALM1-SaCas9-3X HA-WPRE-bGH-U6-gRNA, pX601 plasmid containing APP gRNA sequence was first digested with XhoI/AgeI. The *pCALM1* promoter with flanking homologous sequence overlapping the digested plasmid’s junctions was generated by PCR using the template Addgene plasmid 213973, and then assembled into the digested vector with HiFi DNA Assembly Master Mix (New England Biolabs, E2621). The product was then digested with SacI, and *WPRE* with flanking homologous sequence to the SacI-digested junctions was generated by PCR using template lentiCRISPR v2 and inserted via HiFi assembly. To generate lentiviral SaCas9 CRISPR constructs, pCK002-derived plasmid was linearized by AgeI/PmeI digestion. Two fragments with homologous flanking sequence overlaps were generated to HiFi-assemble into AgeI/PmeI-digested vector: 1) SaCas9-3xHA or SaCas9 KKH-3xHA was amplified by PCR using pX601-derived template; 2) a downstream fragment spanning P2A-Puro-WPRE-3’ LTR was PCR-amplified using pCK002-derived plasmid as template. SpCas9 CRISPR constructs were generated according to lentiCRISPR v2 cloning protocols described in ref # ^67^. Potential off-target effects were assessed using the online tool CRISPROR (http://crispor.tefor.net/ ), and a guide with minimal off-target effects was selected for experiments. All CRISPR sgRNA sequences were analyzed with the National Center for Biotechnology Information’s (NCBI) Basic Local Alignment Search Tool (BLAST) to ensure specificity. AAVs were obtained from the UCSD viral vector core and VectorBuilder. Virus titers of AAVs used in *App*-KI / Cas9 mice were measured by real-time Q-PCR to determine genome copy number of the vector preparations (gc/ml) as a measure of AAV particles with full genome content. Single vector AAV titers were determined by ddPCR (VectorBuilder).

#### AAV injections in vivo

Mice (age: ∼ 5-7-weeks) were anesthetized by isoflurane, and AAVs were injected retro-orbitally using an 8mm, 31G insulin syringes (BD Medical Systems) as described previously ^67, 68^. A dose of 3X10^11 VG/mouse was used for injections in *App*-KI/*Cas9*-KI mice. For single AAV vectors, 4-6-week old *App*-KI mice were treated with an AAV dose of 6x10^13 vg/kg. Mice were maintained 3-10.5 months post-injection for behavioral testing and tissue collection.

### Histology and related procedures

#### General procedures

Mouse hemibrains were quickly dissected, and immediately drop fixed in 4% paraformaldehyde. Tissue was then embedded into paraffin blocks, and 5μm thick sections were collected on Super Frost glass slides (Fisher Scientific). For immunohistochemistry and immunofluorescence, tissue sections were stained with antibodies to appropriate epitopes on a Ventana Discovery Ultra platform (Ventana Medical Systems, Tucson, AZ, USA). Antigen retrieval was performed using CC1 (Tris-EDTA based; pH 8.6) for 40 minutes at 95 ⁰C. Primary antibodies were incubated on the sections for 32 minutes at 37 ⁰C, and were detected using the OmniMap systems (Ventana). Antibody-signal was visualized using DAB as a chromagen followed by hematoxylin as a counterstain. *<u>Multi-color immunofluorescence</u>*: Staining with appropriate antibodies was performed using the OmniMap detection system HRP-coupled secondary antibodies (Ventana) followed by immunofluorescence-labeling according to the manufacturer’s with tyramide signal amplification (TSA)-Alexa 488, TSA-Alexa 647 or TSA-Alexa 594 tyramide-conjugated fluorophores (Invitrogen). To perform multi-color staining, the antibodies were fully denatured, inactivated, and removed from the tissue by treatment in CC2 citrate-based, pH 6.5 (Ventana) for 20 min at 100 °C. Subsequently, the second antibody was applied and detected by the OmniMap system (Ventana) using TSA-Alexa 488, TSA-Alexa 647 or TSA-Alexa 594 tyramide-conjugated fluorophores. After immunofluorescence staining, sections were rinsed and coverslipped with Vectashield containing DAPI (Vectorlabs). Slides were imaged at the UCSD microscopy core using a VS200 (Olympus) or Nanozoomer (Hamamatsu) slide scanner.

#### Quantification of Aβ (6E10), LAMP1 and GFAP staining by immunohistochemistry

Histological sections were scanned at 20x zoom using NanoZoomer slide scanner and the resulting .ndpi files were converted to .tiff images. These high-resolution files were then processed in ImageJ or Qupath. After color deconvolution, the polygonal lasso tool was used to select the anatomical area of interest (i.e. cortex) in DAB channel images and images were converted to 8-bit. After a threshold was generated for each stain, the same threshold was used to process all images within that group. The % area stained was then determined with ImageJ. At least 6-8 tissue sections from each animal were analyzed per stain.

#### Quantification of Aβ (6E10), LAMP1 and GFAP staining by immunohistochemistry

Whole hemispheres were first sectioned to a depth where the full hippocampus could be realized (∼0.8mm from the midline) and at least two consecutive sections for each stain were then taken at this depth. Consecutive sections were then sampled again 40 and 80 sections later. Histological sections were scanned at 20x zoom using NanoZoomer slide scanner and the resulting .ndpi files were converted to .tiff images. These high-resolution files were then processed in ImageJ or Qupath. After color deconvolution, the polygonal lasso tool was used to select the anatomical area of interest (i.e. cortex) in DAB channel images and images were converted to 8-bit. After a threshold was generated for each stain, the same threshold was used to process all images within that group. The % area stained was then determined with ImageJ. At least 6-8 tissue sections from each animal were analyzed per stain.

#### Quantification of activated microglia using the HALO module

Images of Iba-1 stained tissue were analyzed using the Microglial Activation module (IHC images: Version 3.5.3577.214) or Microglia Activation FL module (IF images: Version 3.6.4134) on the HALO Image Analysis Platform (Indica Labs, Inc., Albuquerque, NM).

#### Quantification of editing efficiency (Y188/GFP dual immunofluorescence)

To determine the number of GFP-positive cells positive for APP C-terminus staining in cortices, tissue sections from one-year-old control and APP-gRNA AAV-injected mice were dual-IF stained with APP C-terminus (Y188) and GFP antibodies. For each tissue section, 7 cortical areas were selected and the number of GFP-positive cells positive for APP C-terminus staining were counted. Cells were counted in 4 different tissue sections/mouse.

#### Quantification of transduction efficiency (GFP/NeuN dual immunofluorescence)

To determine the number of NeuN positive cells transduced by AAVs, tissue sections from APP-gRNA AAV-treated mice were dual-IF stained with NeuN and GFP antibodies. GFP positivity was quantified in two dual-labeled sections from each mouse (N = 3 mice). A total of 8 regions were analyzed per section. Briefly, we sampled 3 cortical regions: 1) anterior somatomotor cortex 2) the somatomotor cortex just anterior to the CA2/3 of the hippocampus and 3) the visual cortex just posterior to the hippocampus. Additional samples were made within these regions to capture cortical layers. To quantify NeuN/GFP overlap, images were processed in MetaMorph (Molecular Devices). Images were thresholded to remove background pixels and regions of positive signal were designated. The ratio of NeuN positive regions overlapping with GFP regions was used to determine transduction efficiency.

### Biochemical assays and evaluation

#### Western blots of brain tissue – procedures and analyses

##### Protein isolation

Frozen tissue was homogenized in 0.15 % Triton X-100 with protease inhibitors, pH 7.4 (Pierce) in PBS. The resulting suspension was centrifuged at 21,000 RCF (relative centrifugal force) at 4 °C for 30 minutes, and the resultant supernatant was collected for western blotting.

##### Preparation of WB samples

Protein concentrations of lysates were determined by DC protein assay (BioRad). After protein assays, samples for western-blots were made using 4X LDS buffer (Invitrogen) and reducing agent (Invitrogen) Samples were added to NuPage 4%-12% acrylamide gradient gels (Thermo Fisher) and gel electrophoresis performed using NuPage MES SDS running buffer with antioxidant solution.

##### Blotting

Proteins were transferred to PVDF membranes (0.2um pore) using 20% methanol in Nupage transfer buffer (Thermo Fisher) with antioxidant (Thermo Fisher). Membranes blocked with 5% fat-free milk in PBS and Primary antibodies were diluted in 5% fat-free milk and added to membranes overnight at 4 °C. Membranes were then washed 3X 5minutes with 0.1% Tween 20 in PBS, and incubated in HRP-conjugated secondary antibodies diluted into 5% fat-free milk for 1 hour at room temperature. Membranes were washed again 3x for 5 minutes, incubated in chemiluminescent development solution (Thermo Fisher) and imaged with Chemidoc imager (BioRad).\Western blots were processed and quantified using Image Lab software (version 6.1, BioRad). For quantification, blot images (.scn files) were opened in Image Lab and sample bands delineated using image tools. Band volumes were then quantified by the software.

#### ELISA / MSD quantification of Aβ in mouse brain tissue

For ELISA/Mesoscale Discovery (MSD) assay measurements of Aβ, brains were quickly dissected and snap frozen on dry ice. Frozen brain tissue was then homogenized in 50 mM TBS with protease inhibitors (pH 7.4) and centrifuged at 200,000x g for 22 min at 4 °C. The resulting supernatant was removed, and the remaining pellet was washed once with 50 mM TBS, centrifuged (200,000x g, 22 min at 4 °C) and then solubilized in 6 M GuHCl (Sigma) with protease inhibitors. Samples were sonicated for 30 sec, vortexed at 1800 RPM for 5 minutes, and incubated at 25 °C for 60 min. Samples were centrifuged once more at 200,000x g for 22 min at 25 °C. Finally, the supernatant was collected as GuHCl-soluble fractions. *<u>For ELISAs</u>*: Human Aβ40 and Aβ42 were detected using ELISA kits, according to the manufacturer’s instructions (WAKO 296-64401 for Aβ42, WAKO 298-64601 for Aβ40). Briefly, diluted GuHCl-soluble lysates were added to ELISA plates overnight at 4 °C. Plates were then washed 5x, human Aβ40/42 detection antibodies were added into wells, and plates were incubated for 3 h at 4 °C. After further washing (5x), the stabilized chromogen was added, and incubated for another 30 min at room-temperature in the dark. Reaction was stopped, and absorbance at 450 nm was read using a luminescence microplate reader. *<u>For MSD assays</u>*: Measurements of Aβ38, Aβ40, and Aβ42 were performed on GuHCl-soluble lysates using the V-PLEX Aβ Peptide Panel 1 (6E10) Kit (Cat#K15200E; Meso Scale Discovery). Total Aβ levels were measured using the R-PLEX Human Aβ (Total) Antibody Set (Cat#F218H-3; Meso Scale Discovery).

### Procedures for hippocampal LTP recordings

#### Preparation of hippocampal slices and electrophysiology recordings

Male and female scrambled-gRNA (control) and APP-gRNA injected mice were anesthetized with isoflurane, decapitated, and the brains were processed for acute hippocampal recordings as described previously. ^69^ Briefly, brains were removed and submerged in ice-cold, oxygenated dissection medium containing (in mM): 124 mM NaCl, 3 mM KCl, 1.25 mM KH_2_PO_4_, 5 mM MgSO_4_, 0 mM CaCl_2_, 26 mM NaHCO_3_, and 10 mM glucose. Coronal hippocampal slices were prepared using a Leica vibrating tissue slicer, before being transferred to an interface recording containing preheated artificial cerebrospinal fluid (aCSF) composed of 124 mM NaCl, 3 mM KCl, 1.25 mM KH_2_PO_4_, 1.5 mM MgSO_4_, 2.5 mM CaCl_2_, 26 mM NaHCO_3_, and 10 mM glucose and maintained at 31 ± 10 ⁰C. Slices were continuously perfused with this solution at a rate of 1.75–2 ml/min while the surfaces were exposed to warm, humidified 95 % O_2_/5% CO_2_. Recordings began after at least 2 h of incubation. Field excitatory postsynaptic potentials (fEPSPs) were recorded from CA1b stratum radiatum apical dendrites using a single glass pipette filled with 2 M NaCl (2–3 MΩ) in response to stimulation (twisted nichrome wire, 65 μm diameter) of Schaffer collateral-commissural projections in CA1c stratum radiatum. Pulses were administered at 0.05 Hz using a current that elicited a 50% maximal spike-free response. After establishing a 20-minute stable baseline, LTP was induced by delivering a single episode of 5 ‘theta’ bursts. Each burst consisted of four pulses at 100 Hz, and were separated by 200 ms (i.e., theta burst stimulation or TBS). The stimulation-intensity did not increase during TBS. To ensure that female mice were in the diestrus phase of the estrus cycle at the time of testing, uteri were removed, examined for vascularization and the presence of fluid (estrus phase), patted dry and weighed. ^70^ Mice with uteri in estrus weighing double (∼200-250 mg) that of uteri obtained from mice in diestrus (70-120 mg) were omitted from the study. The fEPSP slope was measured at 10–90% fall of the slope and data in figures on LTP were normalized to the last 20 min of baseline. Electrophysiological measures were analyzed using a one-way ANOVA, and the level of significance was set at p ≤ 0.05.

#### Confirmation of gene-editing in LTP slices

Brain slices were immediately frozen after LTP experiments. Genomic DNA was extracted from frozen slices using QuickExtract buffer (Lucigen). The genomic region containing CRISPR-edits was PCR amplified from genomic DNA template using RedExtract PCR mix (Sigma). Amplicons were purified from PCR buffer using Monarch PCR cleanup kit (New England Biolabs) and submitted for Sanger sequencing (EtonBio).

### Genomic analyses and evaluation

#### Next-generation AMP-sequencing

##### Library preparation and PCRs

The AMP-seq (anchored multiplex PCR sequencing) library preparation protocol was followed as described. ^71^ Briefly, genomic DNA (gDNA) from scrambled-control/*App*-gRNA AAV-injected cortices were used for AMP-seq analysis. For each sample, 30 ng of gDNA was enzymatically fragmented using the KAPA HyperPlus Kit (Roche). Samples were purified at 0.8X using AMPure XP beads (Beckman Coulter), and end-repaired and A-tailed (KAPA HyperPlus Kit). Adapter primers containing amplicons for sequencing, as well as an <u>8-mer barcode</u> and a *random 8-mer* for de-depulication during analysis, were synthesized by Integrated DNA Technologies, annealed, and ligated to fragmented DNA. The first round of PCR was performed using a primer complementary to the adapter and a primer complementary to the genome (5’-TGCCTACGAGTACTGTGCTCC-3’). Following AMPure XP bead cleanup, a second round of PCR was performed using the same adapter primer and a nested primer complementary to an upstream region of the genome . Library concentrations were quantified using the KAPA library quantification kit (Roche), bioanalyzed for quality control (Agilent 2100 Bioanalyzer), and sequenced on an Illumina MiniSeq using the MiniSeq Mid Output Kit with standard sequencing primers (Illumina #FC-420-1002; 150 cycles; paired-end reads). *<u>Data</u> <u>analysis:</u>* Reads were demultiplexed using Je suite^72^ and then filtered such that at least 90% of bases had a Phred quality score ≥ 20 (99.0% accuracy). Reads were then analyzed and binned into editing event categories based on deviation from gRNA reference sequence ± 20 bp from expected Cas9-cleavage site. Reads that were 100% complementary to the reference genome were filtered out as unedited reads. Reads that contained fusion of reference sequence and adapter sequence within ± 20 bp from expected Cas9 cleavage site were also filtered out. We then removed any existing adapter using CUTADAPT and used STAR 2.7.0^73^ (default parameters) to retain only on-target reads that aligned to a 1000 bp window surrounding the gRNA locus. Reads were then aligned to a custom AAV reference genome to identify host genomic reads that were fused to AAV genome within ± 20 bp from expected Cas9 cleavage site. Remaining reads were then re-aligned to the gRNA locus of interest to isolate reads containing insertions or deletions (indels). Substitutions were ignored as presumed sequencing errors, as previously reported. ^74, 75^ Remaining reads were then aligned to an mm10 reference genome (STAR; default parameters) to identify reads containing extra-chromosomal sequence ± 20 bp from the expected Cas9-cleavage site (translocations). All bins were manually spot-checked for accuracy and visualized through CRISPResso2^76^ to confirm accuracy. CRISPResso2 parameters were as follows: CRISPResso -r1 <fastq_file.fastq> -a <amplicon_sequence> -g <grna_sequence> -q 20 -amas 50 --ignore_substitutions -- quantification_window_size 20 --quantification_window_center -3 --exclude_bp_from_left 20 -- exclude_bp_from_right 20 --plot_window_size 40. Data was then quantified as percentage of reads with each editing event category to the total number of reads with any category of editing event.

#### On- and off-target analysis of gene-editing

gDNA from was PCR amplified using primers listed in **Extended Data Table 2**, PCR purified (Qiagen), and Sanger sequenced using Genewiz (Azenta Life Sciences). Data from Cas9- and control-gRNA treated .ab1 files were uploaded to the Synthego ICE CRISPR analysis tool^77^ using default settings to estimate editing frequencies at on- and off-target sites for SpCas9 APP gRNA. Results were plotted as predicted insertion and deletion sizes, as a percentage to total sequences.

#### Next-generation ITR-seq (inverted terminal repeat sequencing

The ITR-seq (inverted terminal repeat sequencing) library preparation protocol was followed as described ^42^. Genomic DNA was extracted from frozen cortex tissue from mice injected with AAV with either an *App*-gRNA or a control-gRNA using DNeasy Blood & Tissue Kit (QIAGEN, no. 69504). gDNA concentration was determined using Qubit 4 (Thermo Fisher Scientific), and the quality was determined using Nanodrop. For each sample (n = 1 each condition), 30 ng of gDNA was enzymatically fragmented using the KAPA Hyperplus Kit (Roche, no. KK8514). The fragmented samples were then end-reparied and A-tailed (KAPA Hyperplus Kit). Custom adapter primers were synthesized by Integrated DNA Technologies. The first adapter primer contains a partial Read 2 sequence and is universal for the samples. The second adapter primer is unique for each sample and contains the P7 sequence, 8 base pair barcode sequence, a random 8-mer for deduplicating reads (UMI), and the Read 2 sequence. The custom adapters were ligated to each other, and then subsequently ligated to the corresponding end-repaired and A-tailed sample fragments. Samples were then purified at 0.8X using AMPure XP beads (Beckman Coulter, no. A63881). A PCR was then performed with a primer complimentary to the adapter sequence containing the P7 sequence, index, random 8-mer, and Read 2 sequence and a primer complementary to the most common ITR breakpoint containing the P5 and Read 1 sequence. The PCR reaction contained KAPA HiFi HotStart ReadyMix (Roche no. 7958927001) and a final primer concentration of 1 μM for each primer. The PCR reaction uses the following amplification conditions: (1) 95C 5min; (2) 98C 20s; (3) 69C 30s; (4) 72C 15s. Repeat steps 2-4 23x. (5) 72C 5min. Samples were then purified with AMPure XP beads at 0.7X. Final library concentrations were measured using Qubit 4 and quality control was assessed with TapeStation (Agilent TapeStation 2200). The pooled libraries were sequenced on an Illumina MiSeq i100 using a 5M Reagent Kit and standard sequencing primers (Illumina, no. 20126565; 2 x 150 bp; paired-end reads with 16 bp index sequences). All primers are listed in Table 1.

#### Preprocessing and ITR-integration analysis

Reads were demultiplexed with 1 mismatched allowed in 8 base pair barcode sequence and deduplicated using the 8 base pair UMI. Reads were then filtered such that at least 90% of bases for each read had a Phred quality score ≥30 (99.9% accuracy). Reads containing a 10 base pair sequence immediately after the ITR primer (ACTCCCTCTC) were filtered as containing potential ITR-genome integration for downstream analysis. These reads were subjected to 5’ trimming of the ITR sequence using CUTADAPT (Parameters: -m 36 -e 0.3) ^78^. The ITR sequence (AGGAACCCCTAGTGATGGAGTTGGCCACTCCCTCTCTGCGCGCTCGCTCGCTCACTGAGGCCGGGCGACCAAAGGTCGCCCGACGCCCGGGCTTTGCCCGGGCGGCCTCAGTGAGCGAGCGAGCGCGCAGCTGCC TGCAGG) was trimmed sequentially in 20 bp fragments, with the remaining sequences undergoing a 3’ trim of 8 Guanines to prevent reads that terminate early from artificially aligning to the mouse genome. Remaining reads were then aligned to an mm39 reference genome using Bowtie2 (Parameters: --sensitive-local) ^79^ with a sensitive local alignment to improve alignment sensitivity and accuracy. Reads were checked by hand to ensure proper alignment to the mouse genome, as some AAV genomic sequences may erroneously align to the mouse genome. The output alignment file was converted to a bedgraph file for visualization in IGV (https://www.nature.com/articles/nbt.1754). Analyses of non-targeting gRNA-treated samples were used as a negative control for ITR integration.

### Behavioral assays and analyses

#### Novel Object Recognition test

Novel object recognition testing was performed as previously described. ^80^ Briefly, on day one, mice were placed into an enclosure with two identical objects. Thereafter, the mice were allowed to explore freely until they reached 20 seconds of total exploration-time of the objects, or 10 minutes total-time in the enclosure, whichever came first. 24 hours later, mice were placed in the same enclosure with one of the objects replaced with a novel object. These mice were free to explore the objects until 20-seconds of total exploration-time, or 10 minutes total-time in the enclosure was reached. The percentage of total exploration-time spent investigating the novel object was calculated.

#### Morris water maze test

A circular pool was filled with ∼23 °C water and made opaque with non-toxic, water-based paint, and the pool was filled until water-level reached 2 cm above the target platform. Visual cues were placed at each quadrant. For each trial of the training phase, mice were placed into the pool and allowed to swim freely for up to 60 sec. Mice that did not find the platform before 60 sec were manually placed onto the platform. For all trials, mice remained on the platform for 20 sec before being removed from the pool. Each mouse completed four trials per day for 8 consecutive days. For probe trial, the target platform was removed on the 9th day of testing. Subsequently, mice were placed into the pool and allowed to swim freely for 60 seconds and the time spent in each quadrant was recorded. Trials were recorded with a webcam (Logitech) and videos analyzed with ANYMaze software. Probe trial data was analyzed with two-way mixed ANOVA with Šídák’s multiple comparisons test to compare time spent in each quadrant between groups. Swim speed and escape latency were analyzed with a 2-way repeated measures ANOVA with Šídák’s multiple comparisons test.

#### Single-nucleus RNA-seq – procedures and analyses

##### Nuclear isolation

Brains from 9-month-old WT, *App-*KI and *App-*KIΔ5 mice were quickly dissected and cut into hemispheres with a single sagittal midline cut, and the hemispheres were flash-frozen and stored at -80 °C. Nuclei were isolated as described in ^81^ with some modifications. Briefly, frozen hemispheres were transferred directly to 5 ml of chilled lysis buffer (10 mM Tris-HCl, 10 mM NaCl, 3 mM MgCl2, and 0.1% Nonidet™ P40 Substitute in Nuclease-Free Water) in a 7 ml glass Dounce homogenizer. Tissues were incubated in the lysis buffer on ice for 5 minutes followed by 3-5 passes with the A pestle, a second 10-minute incubation on ice, and 3-5 passes with the B pestle. Homogenates were then filtered through a 30mm MACs SmartStrainer and centrifuged at 500 rcf for 5 minutes at 4 °C to isolate the nuclear fraction. Nuclear fractions were washed twice with 10 ml of resuspension buffer (1X PBS with 1.0% BSA and 0.2 U/ml Protector RNAse inhibitor (Sigma-Aldrich)). Myelin debris was removed using magnetic separation with Miltenyi Myelin Removal Beads, LS columns and centrifugation. Nuclear pellets were then resuspended in resuspension buffer at a concentration of 1000 nuclei/ ml.

##### Single cell RNA sequencing

Single-nucleus RNA libraries were generated by the UW-Madison Biotechnology Center Gene Expression Center using the Chromium Next GEM Single Cell 3′ Library (10X Chromium). These libraries were subjected to paired end sequencing on the NovaSeq 6000 system (Illumina). Demultiplexed FASTQ files were aligned to the GRCm39 Mus musculus reference genome using Cell Ranger 6.1.2 with the include-introns flag selected. Following alignment, default quality control settings in Cell Ranger were used to filter reads and sort them into cell-associated matrixes. A second round of quality control excluding nuclei with >5% mitochondrial genes, > 50,000 unique molecular identifiers, < 100 or > 7500 genes, as well as all other downstream analyses were performed in Seurat version 4.1.1 (R Studio version 4.2.0).

###### Dimensionality reduction, clustering and cell-type identification

Filtered matrices were log-normalized and highly variable genes were identified for each sample using the FindVariableFeatures function with default parameters. The FindIntegrationAnchors function was then used to identify variable genes conserved across all datasets. These anchors were used to integrate the datasets using the IntegrateData function. The integrated matrix was then scaled and linear dimensionality reduction was performed using the RunPCA function. Next, UMAP and k-nearest neighbor clustering were performed using the functions RunUMAP and FindNeighbors incorporating 20 dimensions, and FindClusters with a resolution of 1. Cell-type identities were assigned to each cluster based on expression of known cell-type markers.

###### Differential Expression Analysis

Cell-type specific transcriptomic profiles from *App*-KI and *App*-KIΔ5 mice were compared to WT by the Wilcoxon rank-sum test using the FindMarkers function with min.pct of 0.1 using Seurat version 5.3.0 (R version 4.3.3). P-values were adjusted using the Benjamini-Hochberg procedure. The level of statistical significance was set at an adjusted p-value < 0.05 and |log2FC| > 0.25.

###### GO analyses

Pathway enrichment analysis was performed using the STRING web tool ^82^. For each cell type, significantly differentially expressed genes were analyzed separately for upregulated and downregulated genes. Genes were included in the enrichment analysis if they met the following thresholds: adjusted p value < 0.05 and absolute log fold change > 0.5. The log fold-change threshold was applied to prioritize genes with a moderate effect size and reduce the contribution of statistically significant but very small expression changes. Enrichment analyses were performed using all STRING-supported pathway annotation databases, excluding PubMed-derived annotations. GO analyses were performed by the Center for Computational Biology and Bioinformatics at UC-San Diego.

##### Screening of gRNAs in iPSC-derived neurons

###### Lentivirus generation

HEK293T cells were maintained in DMEM+GlutaMAX (Gibco, 10566016), 10% FBS (Gibco, A5256701), and 1x penicillin-streptomycin (Gibco, 15140122) at 37°C, 5% CO_2_ and plated on 15 cm dishes coated with 0.1 mg/mL poly-D-lysine (MP Biochemicals, MP215017580). Cells were transfected when 80-90% confluent with 20 ug lentiviral transfer plasmid, 15 ug psPAX2 (Addgene plasmid #12260), and 10 ug pMD2.G (Addgene plasmid #12259) overnight using the ProFection® Mammalian Transfection System (Promega, E1200). Transfection-containing medium was exchanged with fresh medium, followed by 2 virus-containing medium collections at 24 hr and 48 hrs post-media exchange. Collections were pooled and spun at 500 rcf for 15 min at 4°C to remove residual cells. The cleared medium was transferred to a fresh tube and incubated for 4-24 hrs at 4°C with 1x Lenti-X™ Concentrator (Takara, 631232). Precipitated lentivirus was pelleted at 1500 rcf for 45 min at 4°C, then resuspended in 100 ul of 1x HBSS (Gibco, 14025092) per 15 cm dish and stored at -80°C. Lentivirus concentrations were determined using the Lenti-X™ GoStix™ Plus kit (Takara, 631281).

###### Stem Cell culture and NPC generation

Kolf2.1J human induced pluripotent stem cells (hiPSCs; The Jackson Laboratory strain #76500) were maintained under feeder-free condition in StemFlex Medium (Cat#A3349401; Gibco) on hESC qualified matrigel (Cat# 354277;Corning) coated cell culture six-well plates (Cat# 3516; Corning) at 37°C in a humidified incubator with 5% CO₂. Neural progenitor cells (NPCs) were derived from hiPSCs according to a modified protocol of Marchetto et al. (2019). Briefly, hiPSCs were dissociated using Collagenase IV (Cat#17104019; Gibco) and transferred to low-attachment dishes in StemFlex Medium supplemented with 10 μM ROCK inhibitor Y-27632 (Cat#1254; Tocris) for 24 h on an orbital shaker at 90 rpm. Subsequently, the medium was replaced every other day with DMEM/F12 with GlutaMAX™ (Cat#10565018; Gibco) supplemented with B27 minus vitamin A (Cat#12587010; Gibco), N2 supplement (Cat#17502048; Gibco), dand 500 ng/mL Noggin (Cat#HZ-1118; Proteintech) for approximately 7 days to promote embryoid body formation. Embryoid bodies were then plated onto 10-cm dishes coated with poly-L-ornithine (PLO) (Cat#P4957; Sigma-Aldrich) and laminin (Cat#23017015; Thermo Fisher Scientific). After 7 days, neural rosettes were manually collected, dissociated with StemPro™ Accutase™ Cell Dissociation Reagent (Cat#A1110501; Thermo Fisher Scientific), and replated onto fresh PLO/Lam-coated dishes in DMEM/F12 supplemented with N2, B27 minus vitamin A, Noggin, and 20 ng/mL bFGF (Cat#233-FB-500; R&D Systems). Cells were maintained under these conditions for two additional passages to obtain a homogeneous NPC population.

###### NPC culture and neuronal differentiation

KOLF2.1J neural progenitor cells (NPCs) were cultured in DMEM/F12+GlutaMAX supplement (Gibco, 10565042), 1x B27 supplement without vitamin A (Gibco, 12587010), 1x N-2 supplement (Gibco, 17502048), 20 ng/mL basic FGF (R&D Systems, 233-FB), and 1x penicillin-streptomycin (Gibco, 15140122) at 37°C, 5% CO_2_ on Matrigel (Corning, 356234)-coated plates. Passaging was performed using Accutase (Corning, 25-058-CI). NPCs were transduced when 70-80% confluent with lentivirus at 5 MOI for 8-13 hr. Cells were washed 5x with DMEM/F12+GlutaMAX and recovered in complete medium for 2 days. NPCs were then selected for positive transduction with complete medium containing 0.1-0.25 ug/mL puromycin (Invivogen, ant-pr) for 7 days with media exchange every 2-3 days. Selected cells were subsequently maintained in complete medium only. NPC differentiation to cortical neurons was performed primarily as described in ^83^. Culture plates were coated with 5 ug/mL laminin (Gibco, 23017015) for 2-4 hr at room temperature, then with 10 ug/mL poly-L-ornithine (Sigma, P3655) overnight followed by 3x wash with cell culture-grade water. NPCs were seeded onto coated plates at ∼30-40% confluence in NPC medium without penicillin-streptomycin. After overnight incubation and surface attachment, the medium was exchanged to neuronal differentiation medium containing DMEM/F12+GlutaMAX, 1x B27 supplement (Gibco, 17504044), 1x N-2 supplement, 1 ug/mL laminin, 20 ng/mL BDNF (R&D Systems, 248-BDB), 20 ng/mL GDNF (R&D Systems, 212-GD), and 250 ug/mL dibutyryl-cAMP (Selleckchem, S7858). Cells were maintained at 37°C, 5% CO_2_ and media was exchanged every 2-3 days for 21 days before sample collection.

###### Quantification of Aβ peptide levels in neuronal conditioned media

Measurements of Aβ38, Aβ40, and Aβ42 were performed using the V-PLEX Aβ Peptide Panel 1 (6E10) Kit (Cat#K15200E; Meso Scale Discovery). Total Aβ levels were measured using the R-PLEX Human Aβ (Total) Antibody Set (Cat#F218H-3; Meso Scale Discovery).

### Statistics

Unless otherwise specified, data was analyzed with a two-tailed, unpaired t-test or one-way ANOVA with Tukey’s post-hoc test, where appropriate. In all cases, statistical significance was defined as p < 0.05. All data are presented as mean ± SEM.

## Supporting information

All Supp figures

## Acknowledgements

This work was supported by grants to SR from the NINDS (R01AG048218, R21AG052404, UF1NS134063), the Epstein Foundation, the Farmer Family Foundation, and the BrightFocus Foundation (#1042555). Brent Aulston was supported by a postdoctoral fellowship from the Alzheimer’s Association. The work was also supported by the following grants: R35GM119644 (KS), NHGRI T32HG002760 (KG), NINDS R01NS109304 (MJZ), NIA AG076835 (MAW), GM134865 and NS124165 (AA), ACTRI UM1TR005449 (Center for Computational Biology and Bioinformatics), and a grant to the UCSD microscopy core (NINDS P30NS047101). We thank Kathy Krentz and Dustin Rubinstein at the University of Wisconsin Genome Editing and Animal Model core for use of their facilities and services; and Sushmita Roy at the Wisconsin Institute for Discovery for the use of electronic resources. We also thank Atsushi Miyanohara from the UCSD Vector Development Core; Rohan Sharma (UCSD) and Nina Bonaventure (UCLA) for help with image analysis; Brian Head (VA San Diego Health care system/UCSD) for help obtaining mice; and Michael Castle and Albertina Torreblanca Zanca (UCSD) for guidance on AAV design/titer-evaluations. High throughput sequencing was performed by the UNC Integrated Genomics Cores (RRID SCR_022620), which is supported in part by the School of Medicine Office of Research and funding from a Lineberger Cancer Center Support Grant (3P30CA016086-49S1) and the University Cancer Research Fund; and we also thank A. Hepperla (UNC) for technical assistance.

## Author contributions

B.D.A. and S.R. conceived the project and supervised all aspects of the work. BDA performed and analyzed most of the experiments, with technical assistance from D.P.P., K.B.-G., N.C., and N.S. K.G. performed and analyzed the sNuc-Seq experiments under the supervision of K.S. H.O.B. performed the AMP-seq and other genomic assays and data-analyses under the supervision of M.J.Z. E.A.K. performed the hippocampal LTP experiments and analyzed the data under the supervision of M.A.W. L.A.P. designed the AAV vectors and helped in other experiments related to molecular biology. F.S. planned and carried out the experiments in Fig5 and performed the cloning of pCALM1 AAV vectors. S.G.R. assisted with some cell culture experiments and performed MSD analyses. T.R. performed ITR-seq and Amp-seq analysis. Sara R. performed GO analyses and assisted with preparation of RNAseq data. J.S. and A.A. were involved in the generation and initial maintenance of the *App* genomic deletion mice. J.C.-S. helped with the off-target analyses, and T.S. and T.C.S. provided the *App^NL-G-F^* mice.

## REFERENCES

1. Sun, J. & Roy, S. Gene-based therapies for neurodegenerative diseases. Nat Neurosci 24, 297–311 (2021).

2. Frangoul, H. et al. CRISPR-Cas9 Gene Editing for Sickle Cell Disease and beta-Thalassemia. N Engl J Med 384, 252–260 (2021).

3. Gillmore, J.D. et al. CRISPR-Cas9 In Vivo Gene Editing for Transthyretin Amyloidosis. N Engl J Med 385, 493–502 (2021).

4. Wong, C. UK first to approve CRISPR treatment for diseases: what you need to know. Nature 623, 676–677 (2023).

5. Cummings, J. et al. Alzheimer’s disease drug development pipeline: 2023. Alzheimers Dement (N Y) 9, e12385 (2023).

6. van Dyck, C.H. et al. Lecanemab in Early Alzheimer’s Disease. N Engl J Med 388, 9–21 (2023).

7. Honig, L.S. et al. ARIA in patients treated with lecanemab (BAN2401) in a phase 2 study in early Alzheimer’s disease. Alzheimers Dement (N Y) 9, e12377 (2023).

8. Kwart, D. et al. A Large Panel of Isogenic APP and PSEN1 Mutant Human iPSC Neurons Reveals Shared Endosomal Abnormalities Mediated by APP beta-CTFs, Not Abeta. Neuron 104, 1022 (2019).

9. Nixon, R.A. Amyloid precursor protein and endosomal-lysosomal dysfunction in Alzheimer’s disease: inseparable partners in a multifactorial disease. FASEB J 31, 2729–2743 (2017).

10. Small, S.A., Simoes-Spassov, S., Mayeux, R. & Petsko, G.A. Endosomal Traffic Jams Represent a Pathogenic Hub and Therapeutic Target in Alzheimer’s Disease. Trends Neurosci 40, 592–602 (2017).

11. Keeler, C.E. Gene therapy. J Hered 38, 294–298 (1947).

12. Gyorgy, B. et al. CRISPR/Cas9 Mediated Disruption of the Swedish APP Allele as a Therapeutic Approach for Early-Onset Alzheimer’s Disease. Mol Ther Nucleic Acids 11, 429–440 (2018).

13. Konstantinidis, E. et al. CRISPR-Cas9 treatment partially restores amyloid-beta 42/40 in human fibroblasts with the Alzheimer’s disease PSEN 1 M146L mutation. Mol Ther Nucleic Acids 28, 450–461 (2022).

14. Duan, Y. et al. Brain-wide Cas9-mediated cleavage of a gene causing familial Alzheimer’s disease alleviates amyloid-related pathologies in mice. Nat Biomed Eng 6, 168–180 (2022).

15. Holstege, H. et al. Exome sequencing identifies rare damaging variants in ATP8B4 and ABCA1 as risk factors for Alzheimer’s disease. Nat Genet 54, 1786–1794 (2022).

16. Kemaladewi, D.U. et al. A mutation-independent approach for muscular dystrophy via upregulation of a modifier gene. Nature 572, 125–130 (2019).

17. Park, H. et al. In vivo neuronal gene editing via CRISPR-Cas9 amphiphilic nanocomplexes alleviates deficits in mouse models of Alzheimer’s disease. Nat Neurosci 22, 524–528 (2019).

18. Zhang, Y., Chen, H., Li, R., Sterling, K. & Song, W. Amyloid beta-based therapy for Alzheimer’s disease: challenges, successes and future. Signal Transduct Target Ther 8, 248 (2023).

19. Nagata, K. et al. Generation of App knock-in mice reveals deletion mutations protective against Alzheimer’s disease-like pathology. Nat Commun 9, 1800 (2018).

20. Hampel, H. et al. The Amyloid-beta Pathway in Alzheimer’s Disease. Mol Psychiatry 26, 5481–5503 (2021).

21. Fortea, J. et al. Alzheimer’s disease associated with Down syndrome: a genetic form of dementia. Lancet Neurol 20, 930–942 (2021).

22. Jonsson, T. et al. A mutation in APP protects against Alzheimer’s disease and age-related cognitive decline. Nature 488, 96–99.

23. Steubler, V. et al. Loss of all three APP family members during development impairs synaptic function and plasticity, disrupts learning, and causes an autism-like phenotype. EMBO J 40, e107471 (2021).

24. Muller, U.C., Deller, T. & Korte, M. Not just amyloid: physiological functions of the amyloid precursor protein family. Nat Rev Neurosci 18, 281–298 (2017).

25. Maloney, J.A. et al. Molecular mechanisms of Alzheimer disease protection by the A673T allele of amyloid precursor protein. J Biol Chem 289, 30990–31000 (2014).

26. Perez, R.G. et al. Mutagenesis identifies new signals for beta-amyloid precursor protein endocytosis, turnover, and the generation of secreted fragments, including Abeta42. J Biol Chem 274, 18851–18856 (1999).

27. Koo, E.H. & Squazzo, S.L. Evidence that production and release of amyloid beta-protein involves the endocytic pathway. J Biol Chem 269, 17386–17389 (1994).

28. Das, U. et al. Visualizing APP and BACE-1 approximation in neurons yields insight into the amyloidogenic pathway. Nat Neurosci 19, 55–64 (2016).

29. Sun, J. et al. CRISPR/Cas9 editing of APP C-terminus attenuates beta-cleavage and promotes alpha-cleavage. Nat Commun 10, 53 (2019).

30. Ring, S. et al. The secreted beta-amyloid precursor protein ectodomain APPs alpha is sufficient to rescue the anatomical, behavioral, and electrophysiological abnormalities of APP-deficient mice. J Neurosci 27, 7817–7826 (2007).

31. Nordstedt, C., Caporaso, G.L., Thyberg, J., Gandy, S.E. & Greengard, P. Identification of the Alzheimer beta/A4 amyloid precursor protein in clathrin-coated vesicles purified from PC12 cells. J Biol Chem 268, 608–612 (1993).

32. Lai, A., Sisodia, S.S. & Trowbridge, I.S. Characterization of sorting signals in the beta-amyloid precursor protein cytoplasmic domain. J Biol Chem 270, 3565–3573 (1995).

33. Maeder, M.L. et al. Development of a gene-editing approach to restore vision loss in Leber congenital amaurosis type 10. Nat Med 25, 229–233 (2019).

34. Rajendran, L. & Annaert, W. Membrane trafficking pathways in Alzheimer’s disease. Traffic 13, 759–770.

35. Sander, J.D. & Joung, J.K. CRISPR-Cas systems for editing, regulating and targeting genomes. Nat Biotechnol 32, 347–355 (2014).

36. Kurosaki, T. & Maquat, L.E. Nonsense-mediated mRNA decay in humans at a glance. J Cell Sci 129, 461–467 (2016).

37. Chan, K.Y. et al. Engineered AAVs for efficient noninvasive gene delivery to the central and peripheral nervous systems. Nat Neurosci 20, 1172–1179 (2017).

38. Saito, T. et al. Single App knock-in mouse models of Alzheimer’s disease. Nat Neurosci 17, 661–663 (2014).

39. Kreis, A. et al. Overexpression of wild-type human amyloid precursor protein alters GABAergic transmission. Sci Rep 11, 17600 (2021).

40. Torroja, L., Chu, H., Kotovsky, I. & White, K. Neuronal overexpression of APPL, the Drosophila homologue of the amyloid precursor protein (APP), disrupts axonal transport. Curr.Biol. 9, 489–492 (1999).

41. Sasaguri, H. et al. Recent Advances in the Modeling of Alzheimer’s Disease. Front Neurosci 16, 807473 (2022).

42. Breton, C., Clark, P.M., Wang, L., Greig, J.A. & Wilson, J.M. ITR-Seq, a next-generation sequencing assay, identifies genome-wide DNA editing sites in vivo following adeno-associated viral vector-mediated genome editing. BMC Genomics 21, 239 (2020).

43. Griciuc, A. & Tanzi, R.E. The role of innate immune genes in Alzheimer’s disease. Curr Opin Neurol 34, 228–236 (2021).

44. Keren-Shaul, H. et al. A Unique Microglia Type Associated with Restricting Development of Alzheimer’s Disease. Cell 169, 1276–1290 e1217 (2017).

45. Latif-Hernandez, A. et al. The two faces of synaptic failure in App(NL-G-F) knock-in mice. Alzheimers Res Ther 12, 100 (2020).

46. Mehla, J. et al. Age-dependent behavioral and biochemical characterization of single APP knock-in mouse (APP(NL-G-F/NL-G-F)) model of Alzheimer’s disease. Neurobiol Aging 75, 25–37 (2019).

47. Muller, U.C. & Zheng, H. Physiological functions of APP family proteins. Cold Spring Harb Perspect Med 2, a006288 (2012).

48. Chen, W.T. et al. Spatial Transcriptomics and In Situ Sequencing to Study Alzheimer’s Disease. Cell 182, 976–991 e919 (2020).

49. Lacar, B. et al. Nuclear RNA-seq of single neurons reveals molecular signatures of activation. Nat Commun 7, 11022 (2016).

50. Pierce, E.A. et al. Gene Editing for CEP290-Associated Retinal Degeneration. N Engl J Med 390, 1972–1984 (2024).

51. Wang, J. et al. An ultra-compact promoter drives widespread neuronal expression in mouse and monkey brains. Cell Rep 42, 113348 (2023).

52. Kugler, S., Kilic, E. & Bahr, M. Human synapsin 1 gene promoter confers highly neuron-specific long-term transgene expression from an adenoviral vector in the adult rat brain depending on the transduced area. Gene Ther 10, 337–347 (2003).

53. Massaro, G. et al. Systemic AAV9 gene therapy using the synapsin I promoter rescues a mouse model of neuronopathic Gaucher disease but with limited cross-correction potential to astrocytes. Hum Mol Genet 29, 1933–1949 (2020).

54. Loeb, J.E., Cordier, W.S., Harris, M.E., Weitzman, M.D. & Hope, T.J. Enhanced expression of transgenes from adeno-associated virus vectors with the woodchuck hepatitis virus posttranscriptional regulatory element: implications for gene therapy. Hum Gene Ther 10, 2295–2305 (1999).

55. Kleinstiver, B.P. et al. Broadening the targeting range of Staphylococcus aureus CRISPR-Cas9 by modifying PAM recognition. Nat Biotechnol 33, 1293–1298 (2015).

56. Mawuenyega, K.G. et al. Decreased clearance of CNS beta-amyloid in Alzheimer’s disease. Science 330, 1774 (2010).

57. Cruchaga, C. et al. Rare variants in APP, PSEN1 and PSEN2 increase risk for AD in late-onset Alzheimer’s disease families. PLoS One 7, e31039 (2012).

58. Rossner, S., Sastre, M., Bourne, K. & Lichtenthaler, S.F. Transcriptional and translational regulation of BACE1 expression--implications for Alzheimer’s disease. Prog Neurobiol 79, 95–111 (2006).

59. Hick, M. et al. Acute function of secreted amyloid precursor protein fragment APPsalpha in synaptic plasticity. Acta Neuropathol 129, 21–37 (2015).

60. Chavez-Gutierrez, L. et al. The mechanism of gamma-Secretase dysfunction in familial Alzheimer disease. EMBO J 31, 2261–2274 (2012).

61. Devkota, S. et al. Familial Alzheimer mutations stabilize synaptotoxic gamma-secretase-substrate complexes. Cell Rep 43, 113761 (2024).

62. Huang, Q. et al. An AAV capsid reprogrammed to bind human transferrin receptor mediates brain-wide gene delivery. Science 384, 1220–1227 (2024).

63. Moyer, T.C. et al. Highly conserved brain vascular receptor ALPL mediates transport of engineered AAV vectors across the blood-brain barrier. Mol Ther 33, 3902–3916 (2025).

64. Lek, A. et al. Death after High-Dose rAAV9 Gene Therapy in a Patient with Duchenne’s Muscular Dystrophy. N Engl J Med 389, 1203–1210 (2023).

65. Mercuri, E. et al. Onasemnogene abeparvovec gene therapy for symptomatic infantile-onset spinal muscular atrophy type 1 (STR1VE-EU): an open-label, single-arm, multicentre, phase 3 trial. Lancet Neurol 20, 832–841 (2021).

66. Chiou, S.H. et al. Pancreatic cancer modeling using retrograde viral vector delivery and in vivo CRISPR/Cas9-mediated somatic genome editing. Genes Dev 29, 1576–1585 (2015).

67. Sanjana, N.E., Shalem, O. & Zhang, F. Improved vectors and genome-wide libraries for CRISPR screening. Nat Methods 11, 783–784 (2014).

68. Yardeni, T., Eckhaus, M., Morris, H.D., Huizing, M. & Hoogstraten-Miller, S. Retro-orbital injections in mice. Lab Anim (NY) 40, 155–160 (2011).

69. Dong, T.N. et al. Temporal endurance of exercise-induced benefits on hippocampus-dependent memory and synaptic plasticity in female mice. Neurobiol Learn Mem 194, 107658 (2022).

70. Hall, J.A., Cantley, T.C., Galvin, J.M., Day, B.N. & Anthony, R.V. Influence of ovarian steroids on relaxin-induced uterine growth in ovariectomized gilts. Endocrinology 130, 3159–3166 (1992).

71. Zheng, Z. et al. Anchored multiplex PCR for targeted next-generation sequencing. Nat Med 20, 1479–1484 (2014).

72. Girardot, C., Scholtalbers, J., Sauer, S., Su, S.Y. & Furlong, E.E. Je, a versatile suite to handle multiplexed NGS libraries with unique molecular identifiers. BMC Bioinformatics 17, 419 (2016).

73. Dobin, A. et al. STAR: ultrafast universal RNA-seq aligner. Bioinformatics 29, 15–21 (2013).

74. Hanlon, K.S. et al. High levels of AAV vector integration into CRISPR-induced DNA breaks. Nat Commun 10, 4439 (2019).

75. Lackner, M., Helmbrecht, N., Paabo, S. & Riesenberg, S. Detection of unintended on-target effects in CRISPR genome editing by DNA donors carrying diagnostic substitutions. Nucleic Acids Res 51, e26 (2023).

76. Clement, K. et al. CRISPResso2 provides accurate and rapid genome editing sequence analysis. Nat Biotechnol 37, 224–226 (2019).

77. Conant, D. et al. Inference of CRISPR Edits from Sanger Trace Data. CRISPR J 5, 123–130 (2022).

78. Martin, M. Cutadapt removes adapter sequences from high-throughput sequencing reads. EMBnet.journal 17, 10–12 (2011). DOI: 10.14806/ej.17.1.200

79. Langmead, B. & Salzberg, S.L. Fast gapped-read alignment with Bowtie 2. Nat Methods 9, 357–359 (2012).

80. Leger, M. et al. Object recognition test in mice. Nat Protoc 8, 2531–2537 (2013).

81. Krishnaswami, S.R. et al. Using single nuclei for RNA-seq to capture the transcriptome of postmortem neurons. Nat Protoc 11, 499–524 (2016).

82. Szklarczyk, D. et al. The STRING database in 2023: protein-protein association networks and functional enrichment analyses for any sequenced genome of interest. Nucleic Acids Res 51, D638–D646 (2023).

83. Marchetto, M.C. et al. Species-specific maturation profiles of human, chimpanzee and bonobo neural cells. Elife 8 (2019).

