## Supplementary material for "Long term rescue of Alzheimer’s deficits *in vivo* by one-time gene-editing of *App* C-terminus": All Supp figures

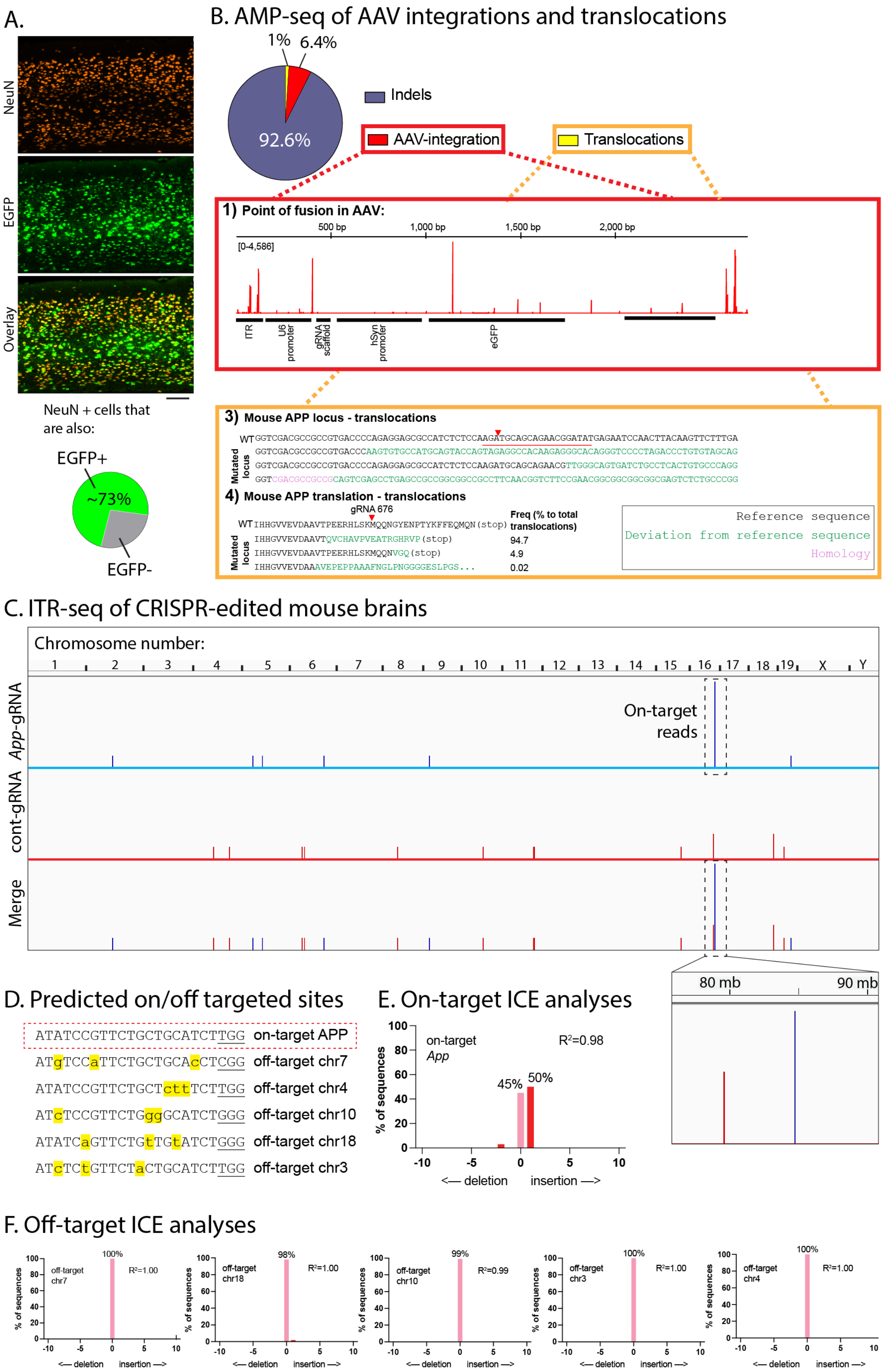


**Extended data Fig. 1: Further characterization of histology and genomics (related to main Fig. 1).**

**A)** Mid-cortical sections from one-year old mouse brains (SpCas9-KI) injected with AAV-PHP.eB-*App*-gRNA (EGFP-tagged), and two-color immunostained for NeuN (neuronal nuclei) and EGFP to determine fraction of transduced neurons. Pie chart on bottom shows the mean number of neurons that were transduced (N = 3 mice per condition, 8 cortical regions and ~ 1000 neurons were analyzed from each animal). These results are consistent with previously published estimates using the AAV-PHP.eB capsid. Scale bar = 100 µm,

**B)** Read pileup demonstrating the point of fusion of the AAV genome with the mouse genome (top, red). The range within the brackets indicates the y axis scale of the number of total reads with AAV integration. Most frequent translocation events (bottom, orange), as evaluated by AMP-seq, with expected translational products as percentage to total translocation events.

**C)** Read pileup demonstrating the genomic location of AAV-ITR integration within the mouse genome, as evaluated by ITR-seq. Tall peak within dotted outline represents on-target editing of the *APP* gene located on mouse chromosome 16.

**D)** Alignment of on-target *App* gRNA site (top row, dashed red box) with predicted off-target sites contain ≤ 3 mismatches (using Cas-OFFinder). All PAM sites are underlined, and nucleotide mismatches in the off-target sites are highlighted in yellow/lowercase.

**E, F).** On-target **(E)** and off-target **(F)** Interference of CRISPR Edits (ICE) analysis of DNA from cortices from *App*-KI/SpCas9-KI mice AAV-injected with *App*-gRNA. Data plotted as frequency of predicted editing outcome (size of insertion or deletion as a percentage to total sequences). Correlation with expected variant percentages (R2) are noted. Note the extremely low frequency of detectable edits within predicted off-target sites. All genomics data from cortex and hippocampus.

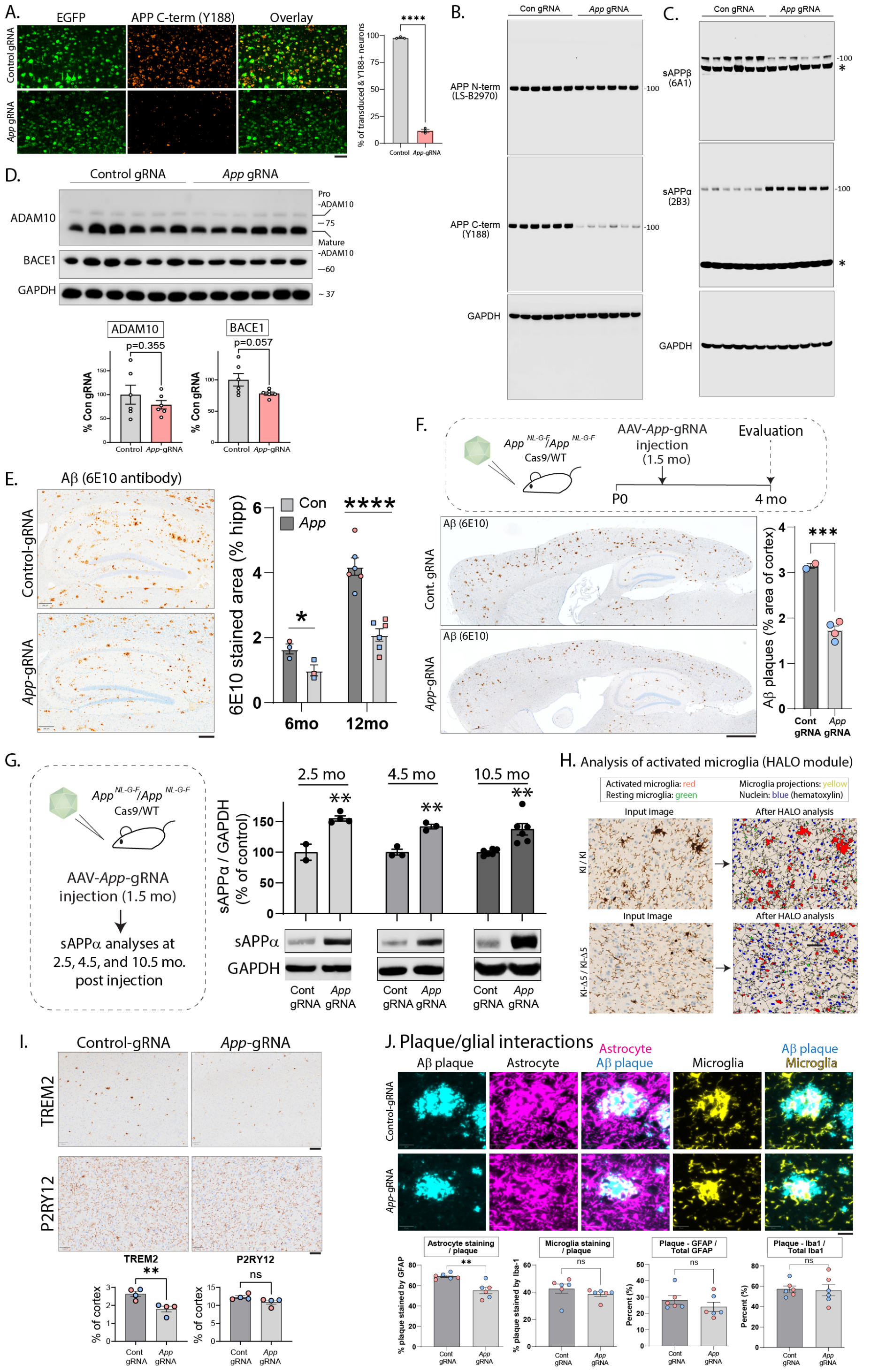


**Extended data Fig. 2: Further characterization and analyses of AAV-driven *in vivo* CRISPR editing (related to main Figs. 1/2).**

**A)** Mid-cortical sections from one-year old SpCas9-KI mouse brains injected with AAV-PHP.eB-*App*-gRNA (EGFP-tagged) or scrambled-gRNA control, and stained for EGFP and Y188 (APP C-terminus antibody) to determine fraction of transduced neurons that also lacked the APP C-terminus; quantified (right). Data shown as mean +/- SEM percentage of EGFP positive neurons that also showed Y188 staining (N = 3 mice per condition, (seven cortical areas were sampled, and >1000 cells were analyzed from each mouse). Note that a large fraction of *App*-gRNA transduced neurons lacked Y188 staining. Scale bar in A) = 50 µm.

**B, C)** Full gels from **Figures 1 H** and **1 I**, asterisk (*) marks non-specific bands.

**D)** One-year old SpCas9-KI mice were injected with AAVs carrying *App*-gRNA or scrambled-gRNA control, and brains were immunoblotted for APP secretases ADAM10 and BACE1; quantified (bottom). Data shown as mean +/- SEM, expressed as a percentage of control-gRNA injected group (N = 6 mice for each condition, ns = non-significant). Note that there are no significant changes in the levels of secretases after *App* C-terminus editing.

**E)** Hippocampal Aβ pathology in APP$^{NL\text{-}G\text{-}F}$/Cas9-KI mouse brains injected with AAV-*App*-gRNA (mean Aβ staining in hippocampi +/- SEM; scale bar = 200µm; red dots in graphs = female mice, blue dots = male mice).

**F-G)** AAV-*App*-gRNA injections attenuated Aβ **(F)** and increased sAPPα **(G)** as early as 2.5 months after injection. Scale bar in **F)** = 1 mm (note that sAPPα blots for 4.5mo and 10.5mo post-injection are crops from **Extended Fig. 3J** and main **Fig. 1I** respectively).

**H)** Automated detection of activated microglia by HALO module software in *App*-KI and *App* -KI-Δ5 tissue sections stained for Iba-1. All western blots were from cortex and hippocampus.

**I)** Immunohistochemistry of DAM (TREM2) and microglial homeostatic (P2RY12) markers in 1yr-old *App*$^{NL\text{-}G\text{-}F}$ / *Cas9*-KI mice injected with *App*-gRNA. Scale bar top = 100 µm, bottom = 50 µm (mean staining per area of cortex, +/- SEM; **p<0.01, ns = non-significant).

**J)** Analysis of glial-plaque interactions in 1yr-old *App*$^{NL\text{-}G\text{-}F}$ / *Cas9*-KI mice injected with AAV-control-gRNA or AAV-*App*-gRNA. Cortical sections were immunostained for Aβ (6E10 antibody – cyan), astrocytes (GFAP antibody – magenta) and microglia (Iba1 antibody - yellow). Representative images are shown with quantification of overlapping plaque–glial staining below. Note that *App* editing led to a significant decrease in the number of astrocytes around plaques, with no changes in other parameters examined (mean +/- SEM, N = 6 animals per group; **p<0.01, ns = non-significant; scale bar = 20µm – unpaired t-test).

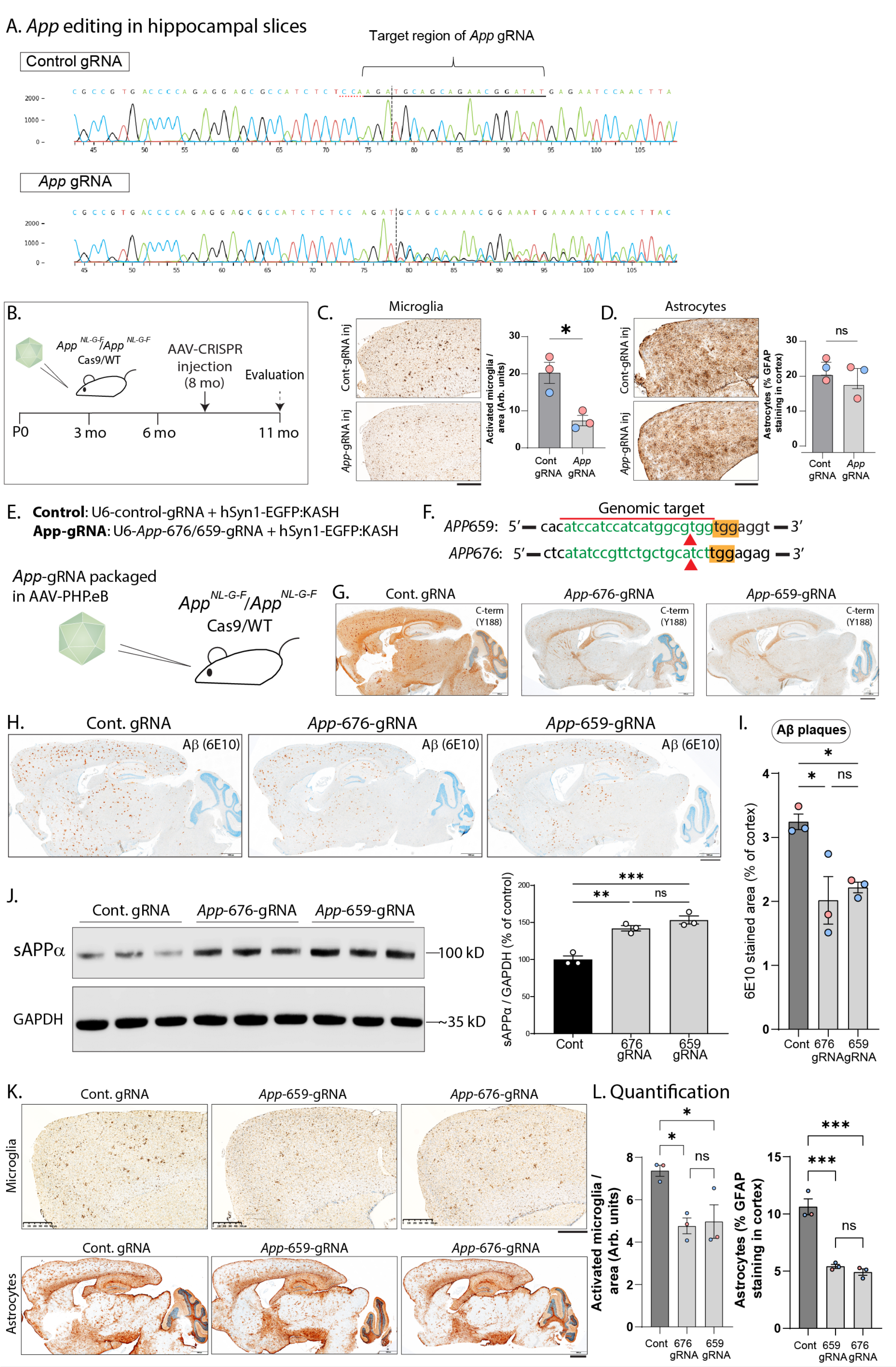


**Extended data Fig. 3**: ***App*-editing in hippocampal slices and comparison of *App*-659 and *App*-676 gRNA (related to main Figs. 2/3).**

**A)** Sanger sequencing confirmation of editing in slices used for LTP experiments. Primers flanking the target site of the *App* gRNA (marked) were designed, and amplicons were produced from the genomic DNA of tissue slices. These amplicons were sequenced to confirm CRISPR-editing. Note nucleotide-mismatch within targeted *App*-gRNA loci, that is seen only in AAV-*App*-gRNA injected brains.

**B-D)** Microglial pathology and astrocytosis in *App*-KI mice injected with AAV-*App*-gRNA at 8 months (after pathology onset, representative frontal cortical sections shown). Note reduction of microglial pathology with a slight (non-significant) decrease in astrocytosis (N=3 mice/group). Scale bars in **C)** and **D)** = 500 µm.

**E)** Experimental design to inject AAV-*App*-659/676 gRNA by intravenous injection in *App*-KI/*Cas9*-KI mice.

**F)** Genomic targets and predicted cut-sites (red arrowheads) in *App* for 659/676gRNAs, PAM sites highlighted yellow.

**G)** Staining of the extreme APP C-terminus (Y188 antibody) in control and 659/676-*App*-gRNA injected mouse brains. Note attenuated staining in *App*-gRNA injected brains, indicating C-terminal truncation of protein. Scale bar = 1 mm.

**H-I)** Attenuated Aβ deposition in 659/676-*App*-gRNA injected mouse brains. Scale bar in **H)** = 1 mm.

**J)** Western blots from mouse brains to examine sAPPα shows an increase in 659/676-gRNA injected mouse brains. All western blots in this figure were from cortical and hippocampal lysates.

**K-L)** Attenuated astrocytosis and microglial activation in 659/676-gRNA injected mouse brains (representative frontal cortical sections), quantified in **L**. Scale bar in = 1 mm for top, 500 µm for bottom images. N=3 mice/group. All data shown as mean +/- SEM, expressed as a percentage of control. N = 3 mice/group, *p<0.05, **p<0.01, ***p <0.001, ns = non-significant (C, D: unpaired t-test, I, J, L: one-way ANOVA).

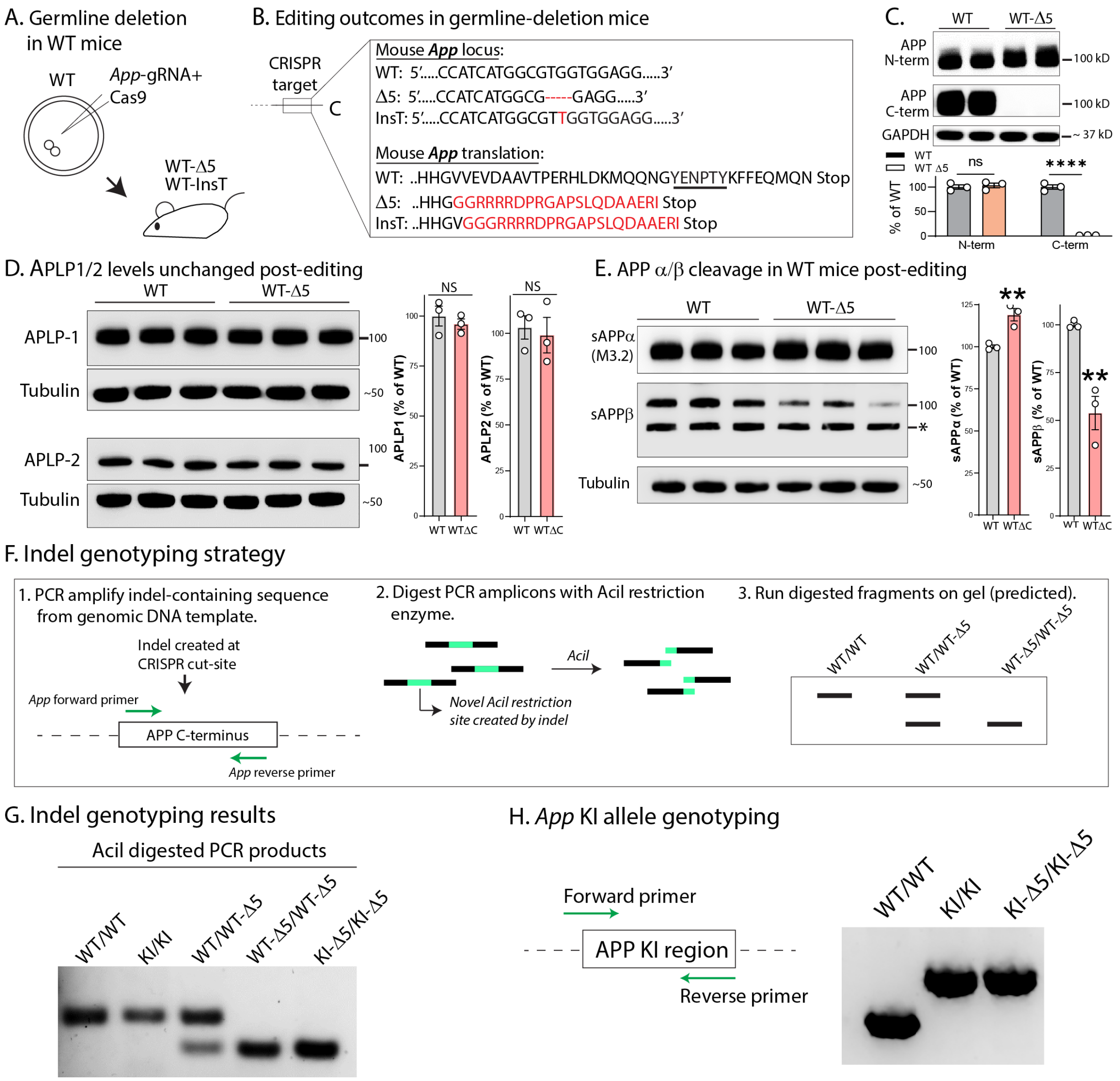


**Extended data Fig. 4: Further characterization of *App*-KI and *App*-KIΔ5/InsT mouse brains (related to main Fig. 3).**

**A)** Zygotes from WT mice were injected with *App*-gRNA/Cas9 ribonucleoprotein complexes targeting the *App* C-terminus, and resulting founders were screened and outbred to produce stable heterozygous and homozygous lines.

**B)** Two edited strains with effective deletion of the last ~35 amino acids were selected. Schematic and sequences show the edited mouse *App* loci and expected translational products. Note that one strain has deletion of 5 amino acids (D5), and the other has a single amino-acid insertion (Insert-T) that lead to indels and premature stop codons.

**C)** Western blots of edited WT mouse brains using antibodies to APP N- and C-terminus. Note that the C-terminus Y188 antibody is unable to recognize APP, while signal from the N-terminus antibody is intact.

**D)** Western blots for APLP1 and APLP2 levels (APP homologues) from WT and WT-Δ5 mice; quantified on right. Data expressed as mean +/- SEM, expressed as a percentage of WT. N = 3 mice per condition, NS = non-significant (unpaired t-test).

**E)** Western blots for sAPPα and sAPPβ from WT and WT-Δ5 mice. Asterisk (*) indicates a non-specific band. Data shown as mean +/- SEM, expressed as a percentage of WT. N = 3 mice/group, **p <0.001 (unpaired t-test).

**F)** Genotyping strategy to identify Δ5bp indel in *App* genomic deletion mice. Deletion of the 5bp generates a novel restriction-site that was digested by the restriction-enzyme AciI, allowing us to identify the genomically deleted mice. Specifically, indel-containing sequences were PCR-amplified, digested with AciI, and the digested fragments were run on agarose gels to identify genomically deleted animals.

**G)** DNA gel of PCR products from WT and *App*-KI mice after digestion with AciI. Note that genomic deletion of the *App* C-terminus (right 3 lanes) generates a cleavage product that runs as a lower band.

**H)** Genotyping to confirm retention of the *App* knock-in allele in KI-Δ5bp mice. Note that presence of the *App*-KI allele leads to an upper band, allowing identification of KI animals. All western blots were from cortical/hippocampal lysates. DNA for genomic analyses was from clipped ear tissue.

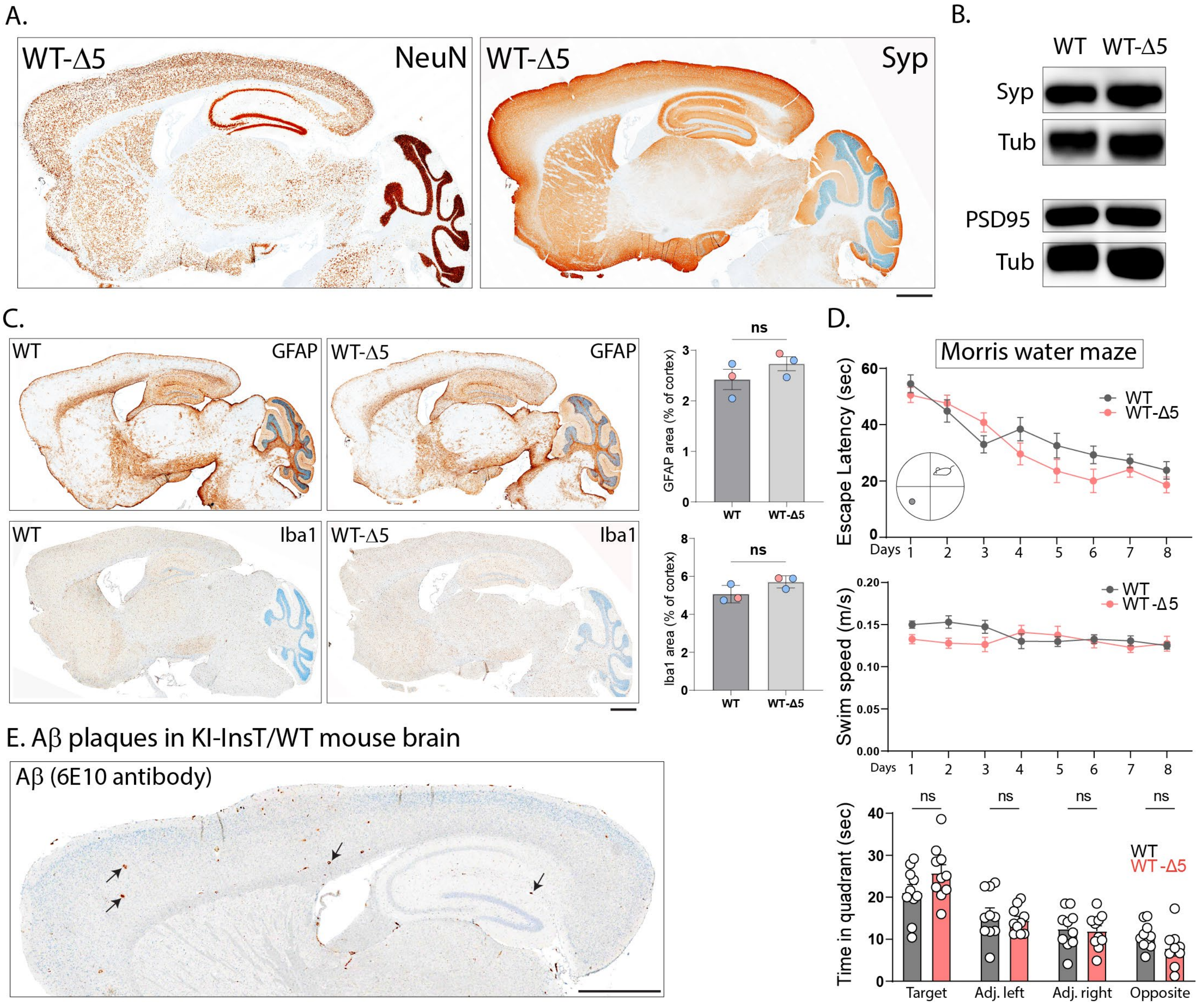


**Extended data Fig. 5: Evaluation of *App*-editing in WT mice (related to main Fig. 3).**

**A-B)** Neuronal (NeuN) and synaptic (synaptophysin) staining in the WT germline-edited mice (*App*-Δ5, a strain that effectively lacks the YENPTY-motif). Note that the brains appear grossly normal. Immunoblots of pre- and post-synaptic markers (whole brain lysates) in **B)**. Scale bar = 1mm.

**C)** Comparison of astrocytes and microglia show no significant changes in the WT germline-edited mouse brains, compared to WT controls; quantified on right. Scale bar = 1mm. Red dots in graphs = female mice, blue dots = male mice (N=3 mice per condition, mean +/- SEM, ns = non-significant, unpaired t-test). Scale bar = 1mm.

**D)** Morris water maze test for memory does not show any deficits in WT germline-edited mouse brains, compared to WT controls (escape latency – top, swim speed – middle, probe-trial results – bottom). N=10 mice/condition, mean +/- SEM, ns = non-significant – two-way ANOVA.

**E)** Representative image of Aβ (6E10) staining in *App*-KI-InsT/WT heterozygous mice lacking the last ~ 35 amino acids of APP. Note that plaques are rarely seen in this setting (few marked with small arrows). Quantification of Aβ in these mice is shown in **Figure 3C**. Scale bar = 1mm

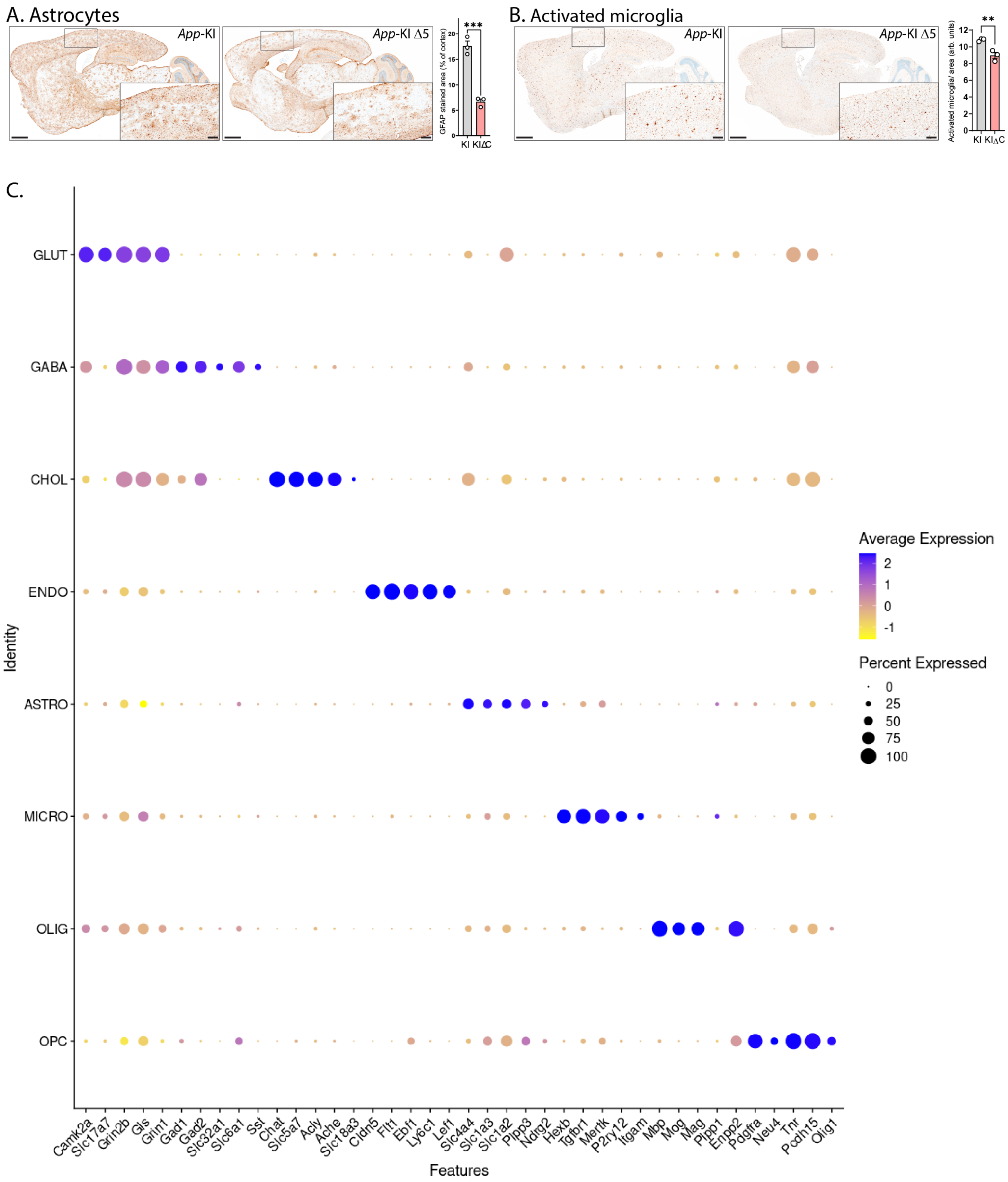


**Extended data Fig. 6: Dot-plots for cell signatures relevant for sNucSeq experiments (related to main Fig. 3).**

**A, B)** Representative histologic sections showing that astrocytic and microglial activation in *App*-KI mice (homozygous) is ameliorated by germline editing of the *App* C-terminus; staining quantified on right (N = 3 animals per condition, mean +/- SEM, **p <0.01, ***p <0.001 – unpaired t-test). Scale bars: main = 1 mm, zoomed inset = 200 μm.

**C)** Dot-plot representation of marker gene-expression for the cell types that were identified. The size of the dot represents the percentage of nuclei in the cluster expressing the marker gene, and the color indicates the expression level relative to the average expression across all nuclei (the data is scaled such that the average expression across all nuclei = 0). Yellow indicates lower than average expression; a graduated shift in color from yellow to purple to dark blue indicates increasing levels of expression with dark blue indicating the highest level of expression. Note that expression of marker genes is largely restricted to the appropriate cell-types, indicating successful cell-type clustering and identification of cell-types.

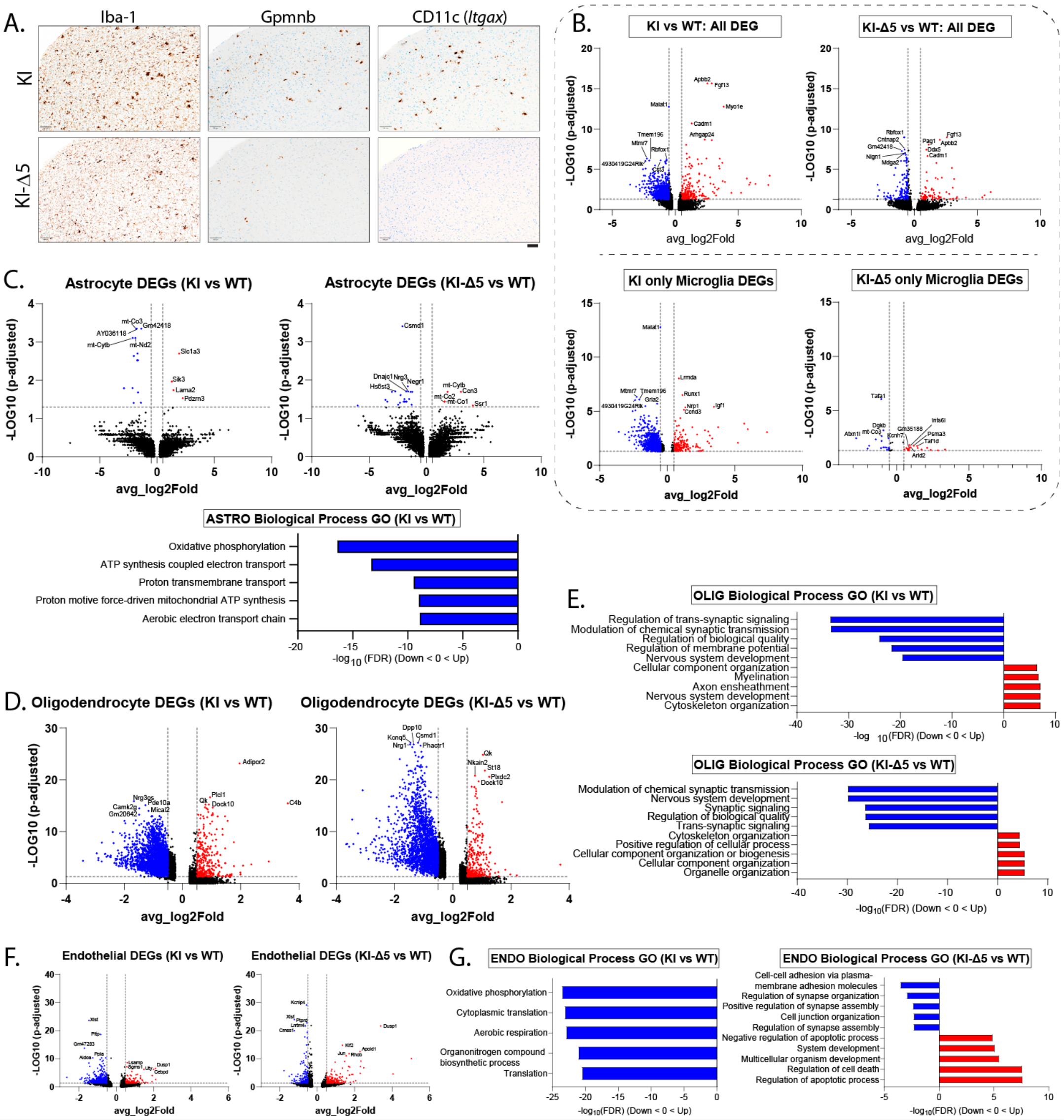


**Extended data Fig. 7: Further analyses of the Snuc-RNA-seq dataset (related to main Fig. 3).**

**A)** Immunohistochemistry showing decreased expression of DAM markers Gpnmb and CD11c (*Itgax*) in *App*-KI/KI-Δ5 brains (cortices). Iba1 staining shows expected decrease of activated microglia in *App*-KI animals after gene-editing. Scale bar = 100 µm.

**B)** Top: Volcano plots of all differentially expressed genes (DEGs) in KI/KI-Δ5 microglia, compared to WT. Colored dots (blue = down-regulated, red = up-regulated) show DEGs with an adjusted p-value of <0.05 and average log2Fold change in expression >0.5. Black dots show genes with a p-value >0.05 and/or log2Fold change in expression of <0.5. Top five significantly up-regulated/down-regulated DEGs are labeled in each graph. Bottom*:* Volcano plots of genotype-specific DEGs in KI/KI-Δ5 microglia, compared to WT. Note substantial attenuation of all DEGs and microglial DEGs upon *App* editing.

**C)** Top: Volcano plots of all DEGs in KI/KI-Δ5 astrocytes, compared to WT. Note attenuation of astrocytic DEGs upon *App* editing. Bottom: Biological Process gene-ontology (GO) analysis of astrocyte DEGs. Bars indicate the top five significantly dysregulated processes in KI astrocytes, compared to WT. Note that no bar-graph is shown for KI-Δ5 astrocytes compared to WT because no biological process GO pathways were significantly enriched in this group.

**D-E)** Volcano plots of all DEGs in KI/KI-Δ5 oligodendrocytes, compared to WT **(D)**, with biological Process GO analysis of oligodendrocytes DEGs in **(E)**. Bars indicate the top five significantly down-regulated (blue) or up-regulated (red) processes in KI/KI-Δ5 (vs WT) oligodendrocytes.

**F-G)** Volcano plots of all DEGs in KI/KI-Δ5 endothelial cells, compared to WT **(F)**, with biological Process GO analysis of endothelial DEGs in **(G)**. Bars indicate the top five significantly down-regulated (blue) or up-regulated (red) processes in KI/KI-Δ5 (vs WT) endothelial cells.

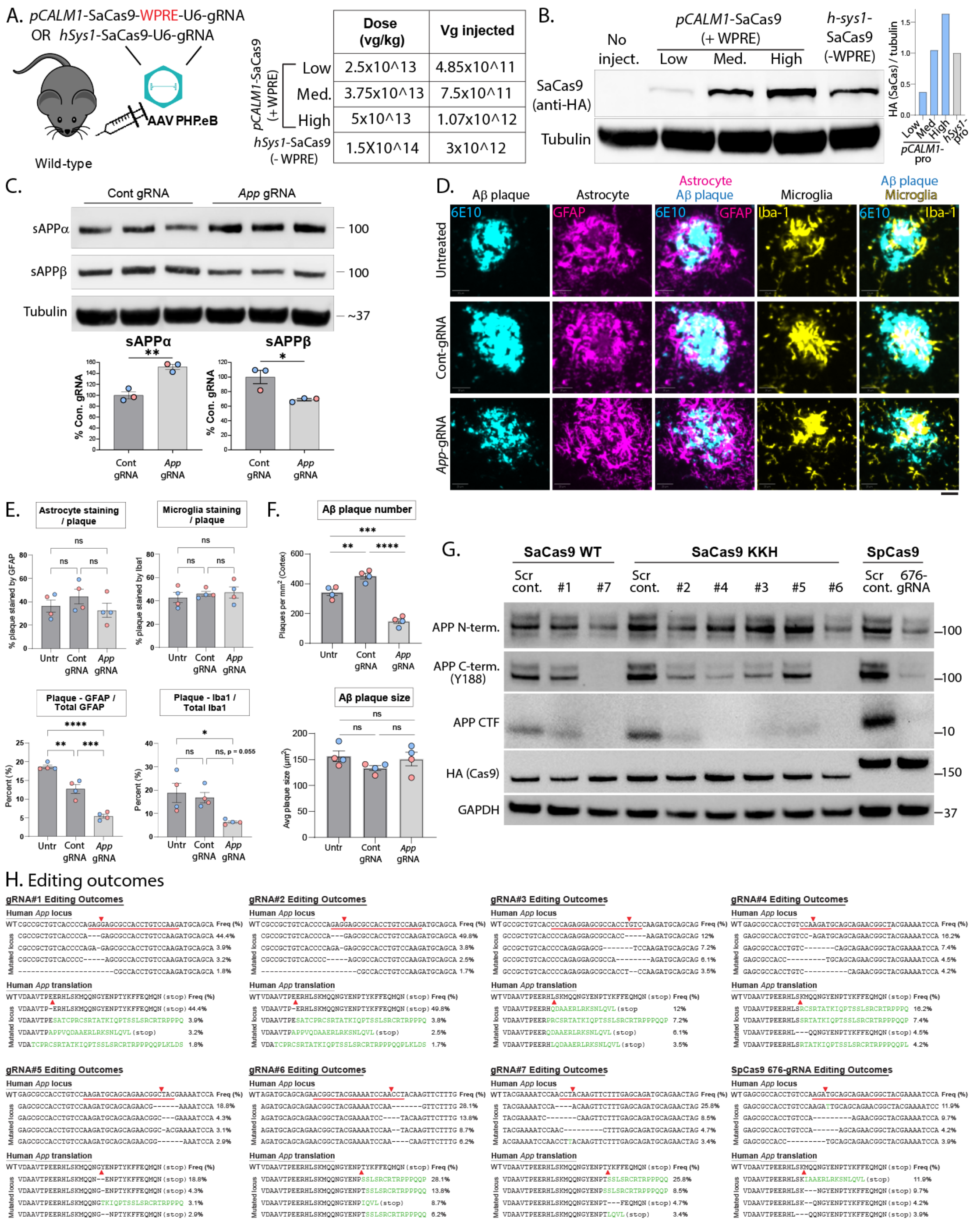


**Extended data Fig. 8: Further characterization of single-AAV vector, glial-plaque interactions, and human APP editing (related to Figs. 4-5).**

**A-B)** In vivo expression (using AAV-PhP.eB) of SaCas9 driven by the *pCALM1* promoter (with *WPRE* element) or human synapsin-1 promoter (no *WPRE* element); table shows injected doses. Mice were injected at 7 weeks, and brains were examined after 4 weeks. Westerns blots from brain lysates (cortex/hippocampus) in **(B)** show HA-tagged Cas9 expression.

**C)** Western blots for sAPPα/sAPPβ from brains of *App*-KI mice injected with a single AAV carrying all CRISPR-components to edit *App* (or control-gRNA – injected at 1.5 months, evaluated at 4 months); quantified below (red dots denote female mice, blue dots male mice). Note increase in α-cleavage and decrease in β-cleavage upon APP editing (mean +/- SEM, N=3 animals per group. *p<0.05, **p<0.01, unpaired t-test).

**D)** Analysis of glial-plaque interactions in single AAV treated brains. Brain sections were stained for Aβ plaques (6E10 antibody – cyan), astrocytes (GFAP antibody – magenta) and microglia (Iba1 antibody - yellow). Scale bar = 20µm.

**E)** Quantification of plaque–glia stain overlap. Though *App* editing did not lead to significant changes in the percentage of glial cells surrounding individual plaques (top panels), there was an overall reduction in the ratio of both astrocytes and microglia around plaques to total glia in the *App* edited groups (bottom panels), likely due to an overall decrease in neuroinflammation in this cohort (mean +/- SEM, N=4 animals per group; *p<0.05, **p<0.01, ***p<0.001, ****p<0.0001; red dots denote female mice, blue dots male mice, one-way ANOVA).

**F)** Quantification of Aβ plaque number per area (top) and mean Aβ plaque size (bottom) in cortices from single-AAV injections (mean +/- SEM, N=4 animals per group; ** p<0.01, *** p<0.001, ****p<0.0001; red dots denote female mice, blue dots male mice, one-way ANOVA).

**G)** Western blots for APP in lentiviral-transduced, puromycin-selected NPC lysates.

**H)** Amp-seq analysis of indels produced at gRNA target sites with expected translational products (four most common outcomes are shown). Red underlines represent gRNA target sequences and red triangles mark expected CRISPR cut-sites. Missense amino acids as a result of editing are shown in green.
